# Embodied Cognition and Decision-Making in the Primate Superior Colliculus

**DOI:** 10.1101/2020.08.14.251777

**Authors:** E.J. Jun, A. Bautista, M.D. Nunez, T. Tak, E. Alvarez, M.A. Basso

**Author notes:** Correspondence and request for materials should be made to M.A.B. These authors contributed equally to this work (E.J. Jun, A. Bautista, M.D. Nunez).

## Abstract

A popular model of decision-making suggests that in primates, including humans, decisions evolve within forebrain structures responsible for preparing voluntary actions; a concert referred to as embodied cognition. Embodied cognition posits that in decision tasks, neuronal activity generally associated with preparing an action, actually reflects the accumulation of evidence for a particular decision. Testing the embodied cognition model causally is challenging because dissociating the evolution of a decision from preparing a motor act is difficult, if the same neuronal activity instantiates both processes. Ideally, one would show that manipulation of neuronal activity thought to be involved in movement preparation actually alters decisions, and not movement preparation. Here, trained monkeys performed a two-choice perceptual decision-making task in which they judged the orientation of a dynamic Glass pattern before and after unilateral, reversible inactivation of a brainstem area involved in preparing eye movements, the superior colliculus (SC). Surprisingly, we found that unilateral SC inactivation produced significant decision biases and changes in reaction times consistent with a role for the SC in evidence accumulation. Fitting signal detection theory and sequential sampling models (drift-diffusion and urgency-gating) to the data revealed that SC inactivation produced a decrease in the relative evidence for contralateral decisions. Control experiments showed that SC inactivation did not result in eye movement biases ruling out interpretations based on motor preparation or spatial attentional impairment. The results provide causal evidence for an embodied cognition model of perceptual decision-making and provide compelling evidence that the SC of primates plays a causal role in how evidence is accumulated for perceptual decisions, a process that is usually attributed to the cerebral cortex.

---

Trained monkeys (Macaca mulatta) perform a two choice perceptual decision-making task in which they viewed a dynamic Glass pattern stimulus^1,2^ and reported the perceived orientation of the stimulus by making saccades to a target located in either the left or right hemifield, before and after reversible, unilateral inactivation of the SC (Fig. 1). Monkeys’ decisions occurred in two conditions: one with the cue to report the choice occurring with a delay after the appearance of the Glass pattern (Fig. 1a; Delay task) and a second with no delay, allowing monkeys to report their decisions immediately (Fig. 1b; RT task). The latter version, known as a reaction time (RT) task, allowed us to fit dynamic decision-making models to the behavioral data to determine what decision-making processes changed, if any, after unilateral muscimol inactivation of the SC.

**Fig. 1.**
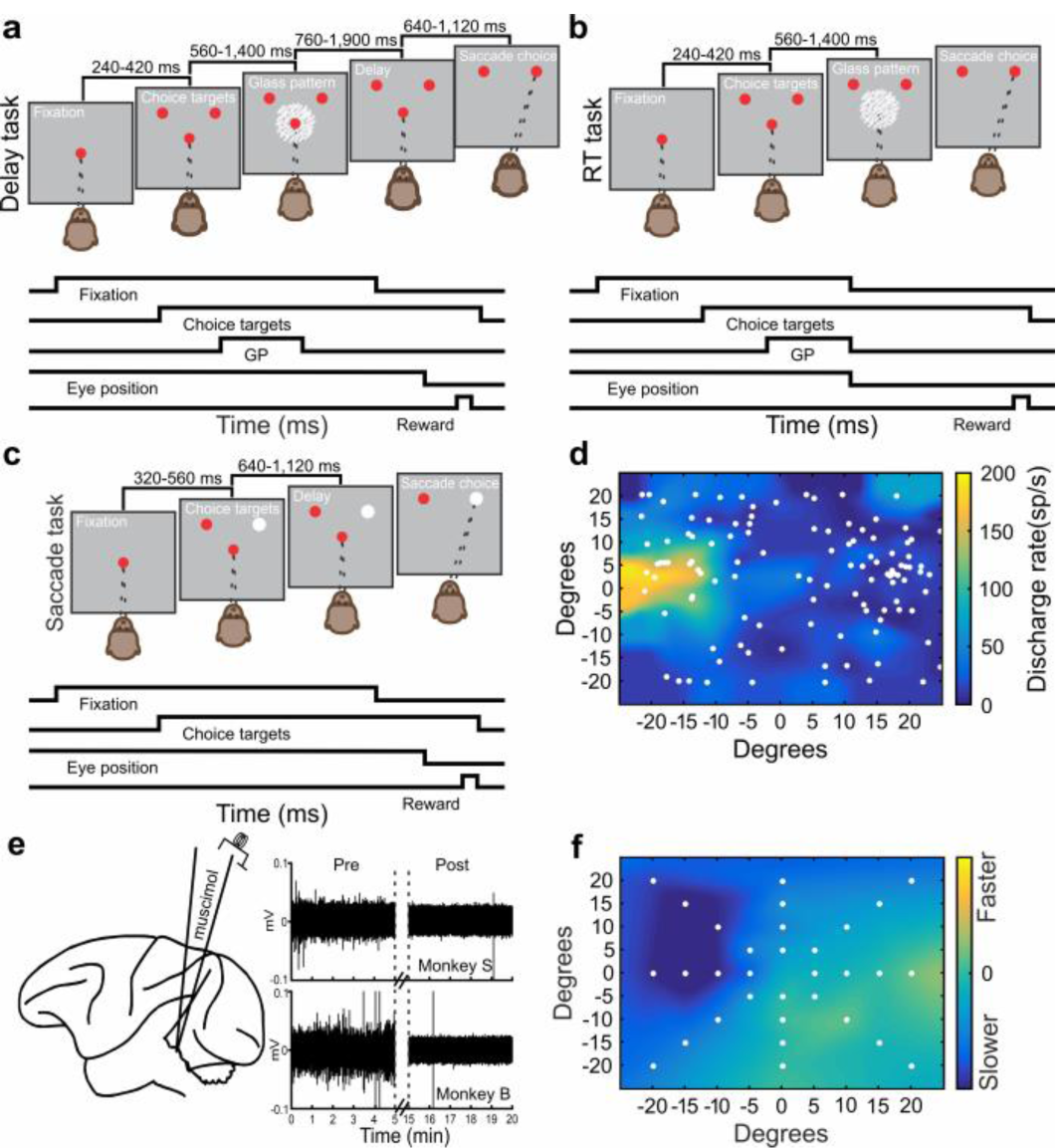
Decision and selection tasks before and after unilateral SC inactivation. **a** The spatial arrangement of the Glass pattern decision task (Delay Task) appears as a sequence of grey boxes depicting the visual display and the temporal sequence of the events appears as running line displays below the spatial display. **b** Same as in a for the reaction time version of the task (RT Task). **c** Schematic display of the saccade selection task in which monkeys made saccades to a white stimulus which varied between the two locations randomly on each trial with a red stimulus. The timings of the tasks appear above the schematic illustrations. **d** The pre-muscimol discharge rate of an SC neuronal recording measured 50ms before and 50ms after the onset of a saccade, is plotted as a heat map using linear interpolation between the target positions (white circles). Warmer colors indicate higher discharge rates (sp/sec). The position 0 horizontal and 0 vertical in degrees marks the position of the centrally-located fixation spot. **e** Schematic of the experimental arrangement showing a lateral view of the rhesus monkey brain and an injectrode targeting the SC. The traces to the right show the raw voltage traces against time in min before and after muscimol injections in two monkeys (monkey S; top and monkey B; bottom). The 10 min injection time is marked by the vertical dashed lines. **f** Post-muscimol saccade velocity minus pre-muscimol saccade velocity is plotted for the target positions indicated by the white circles. Cooler colors indicate slower saccadic velocities post-muscimol. The plots shown in d and f were taken from one injection experiment in monkey S. The location of the response field (RF) as in d, determined the positions of the choice targets in the decision and selection tasks.

Monkeys also performed a saccade selection task in which two possible choice targets appeared at the same two locations as in the decision task. One target was red and the other white, and the monkeys made saccades to the white target, which alternated between the left and right positions randomly on each trial (Fig. 1c). The selection task required the same attention to the peripheral location and the same motor preparation as in the decision task but did not require the transformation of an orientation decision to a motor action. Performing this task before and after unilateral inactivation of the SC assessed motor preparation and bias.

Fig. 1d shows a heat map of neuronal discharge recorded from an SC neuron before a muscimol injection. The warmer colors in Fig. 1d show a typical SC neuronal response field (RF) measured while a monkey made visually-guided saccades to the locations shown by the white circles. We positioned the choice targets for the decision and selection tasks at the RF center (to inactivated field (toIF)) and 90° opposite (awayIF). Muscimol reliably reduced or silenced the spontaneous activity of SC neurons within 10-15 minutes of the injection (Fig. 1E). Monkeys also performed visually-guided saccades to predefined targets before and after inactivation, allowing us to map changes in saccadic velocity and providing an independent, behavioral measure of the efficacy of the muscimol injection^3-6^ (Fig. 1f). Extended Data Table 1 shows details of the injections made in two monkeys.

## SC inactivation biases decision-making

Unilateral muscimol injection into the SC produced a reliable ipsilateral (awayIF) decision bias in both monkeys for all 23 injections in both the delay and RT tasks (Fig. 2a). Unilateral saline injections into the SC (n=6) produced little discernible effect on decision-making performance (Fig. 2b). Parameter estimation from logistic fits to the performance data revealed a statistically significant rightward shift in *α*, the decision bias parameter (Fig 2c, pre *α* = -0.29, post *α* = 10.21, t(22) = -9.81, p = 1.725 × 10^−9^). In some experiments, the perceptual sensitivity parameter *β*, decreased with muscimol, but on average the decrease failed to reach significance with Bonferonni correction (Fig 2d, pre *β* = 0.12, post *β* = 0.10, t(22) = 2.23, p = 0.037). Saline injections affected neither parameter (Fig. 2e, f; pre *α* = 0.13, post *α* = 0.07, w(5) = 14, p = 0.563; pre *β* = 0.12, post *β* = 0.11, t(5) = -1.71, p = 0.148). Calculating signal detection theory (STD) quantities, *d’* and criterion (*c*) also revealed statistically significant changes in *c* and not *d’* (Extended Data Fig. 1).

**Fig. 2.**
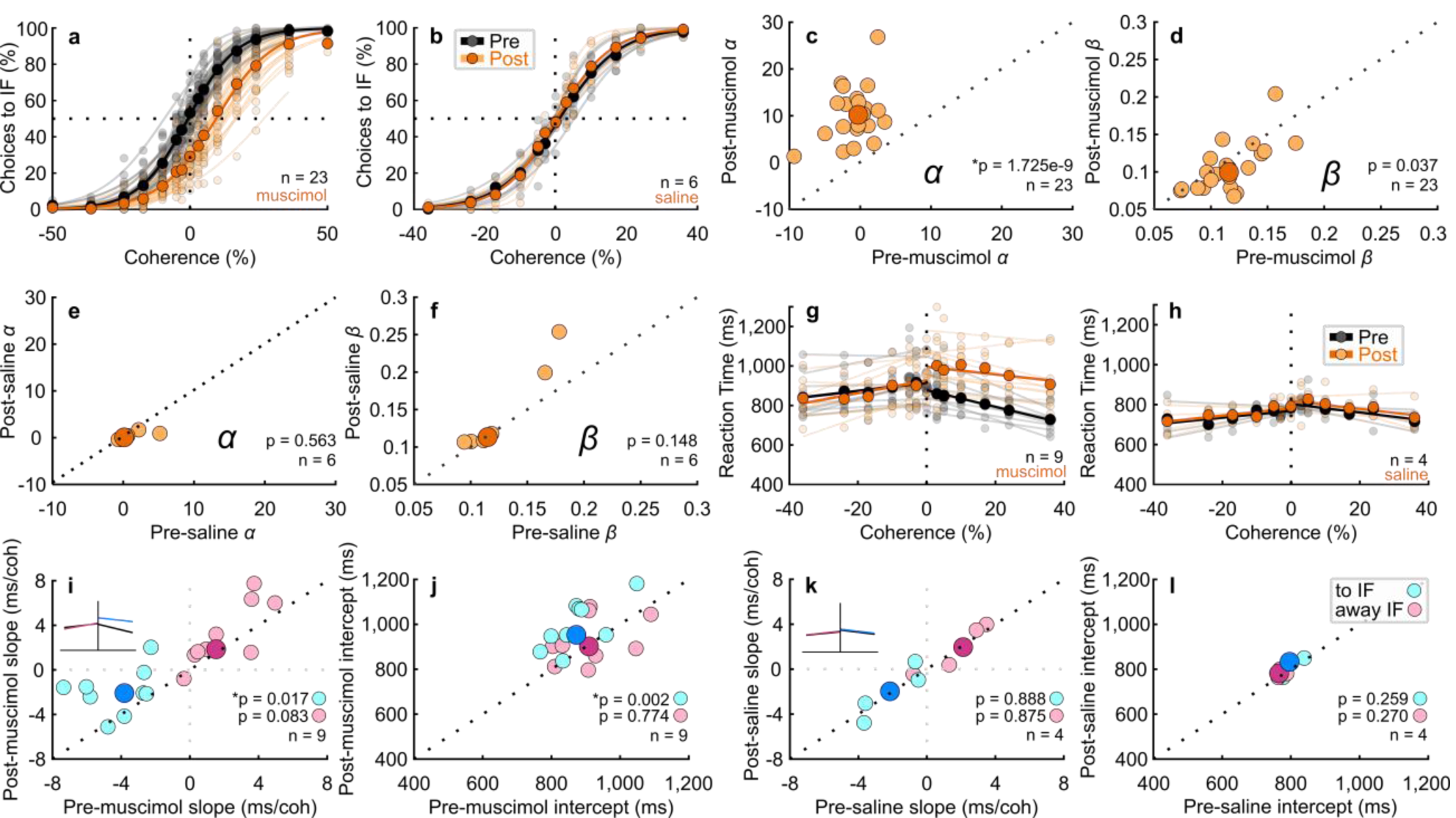
Unilateral Inactivation of the SC biases perceptual decision making. **a** Decisions to the inactivated field (toIF) plotted against Glass pattern coherence, where positive coherences presented toIF evidence and negative coherences presented awayIF evidence, for 23 experiments performed in two monkeys before (black circles and lines) and after (orange circles and lines) unilateral injection of muscimol into the SC; n = 11 in monkey B and n = 12 in monkey S for both delay and RT tasks. The horizontal dashed lines show 50% chance performance. Vertical dashed lines show 0% coherence. Each lighter shade line shows the two parameter logistic fit to the data for individual experiments and the darker shade lines show the two parameter logistic fit pooled across 23 experiments (see Extended Data Fig. 2 for comparisons of two, three and four parameter logistic fits). Extended Data Fig. 3 shows the results of the 24-hour recovery for muscimol and saline and Extended Data Table 2 shows the associated statistics. **b** Same as in a for the Glass pattern task performance before (black) and after (orange) saline injections, n = 2 in monkey B and n = 4 in monkey S. Note that there are no 50% coherence for saline experiments (Methods). **c-f** Post-muscimol and post-saline parameters of the logistic fits are plotted against the same parameters measured pre-muscimol and pre-saline from the fits shown in a and b. The darker symbols show the medians whereas the lighter symbols show the parameters from individual experiments. * indicates significance with a critical *α* value of 0.025 Bonferroni corrected. **g** For the same experiments and data shown in a that were performed with the RT version of the decision task (n=9), RT is plotted against coherence pre-muscimol (black circles and lines) and post-muscimol (orange circles and lines) for all correct trials. The lines show linear fits to the data points (Supplementary methods). **h** The same as in g for the saline injections (n=4). **i** The slopes of the post-muscimol linear fits to the RT data are plotted against the pre-muscimol slopes for toIF decisions (cyan circles) and awayIF decisions (magenta circles). The inset shows which changes in slope correspond to the changes seen in the chronometric function. The black dotted line shows unity. The dark circles show medians. **j** Same as in i for the intercept parameter. **k-l** The same as in i-j for the saline experiments.

Of the 23 muscimol injections, nine were performed during the RT version of the task (seven in monkey S and two in monkey B). On average, mean RT increased for toIF decisions and did not change for awayIF decisions post-muscimol (Fig. 2g; mean RT pre toIF = 808.71 ms, mean RT post toIF = 971.77 ms, *t*(53) = -12.86, p = 6.24 × 10^−18^; mean RT pre awayIF = 887.31 ms, mean RT post awayIF = 871.41 ms, *t*(53) = 1.11, p = 0.27). However, mean RTs showed a negative correlation with coherence for toIF decisions before the injection and the slopes of the chronometric functions flattened after injections (Fig. 2g, Fig. 2i cyan circles, pre = -3.83 ms/coherence, post = -2.09 ms/coherence, *t*(8)= -3.00, p = 0.02). We found no significant changes in slopes for awayIF decisions (Fig. 2g, Fig. 2i magenta circles, pre = 1.55 ms/coherence, post = 1.84 ms/coherence, *t*(8) = -1.98, p = 0.08). Significant changes occurred in the intercept parameter for toIF decisions but not for awayIF decisions (Fig. 2j cyan circles, pre toIF = 873.085 ms, post toIF = 952.62 ms, *t*(8) = -4.64, p = 0.002; magenta circles, pre awayIF = 909.725 ms, post awayIF = 901.225 ms, *t*(8) = -0.30, p = 0.77). The significant changes in slope of the chronometric functions for toIF decisions indicate a unilateral, coherence-dependent change in mean RT. That the lower coherence trials show less of a change in RT than the higher coherence trials, indicates that the change in RT is not solely a result of motor impairment.

The four saline injections changed neither the slope nor intercept of the chronometric functions (Fig 2h; Fig. 2k cyan circles, pre slope toIF = -2.14 ms/coherence and post slope toIF = -1.99 ms/coherence, *t*(3) = -0.15, p = 0.89; magenta circles, pre slope awayIF = 2.11 ms/coherence and post slope awayIF = 1.94 ms/coherence, *w*(3) = 4.00, p = 0.88; Fig. 2l cyan circles, pre intercept toIF = 796.70 ms and post intercept toIF = 832.71 ms, *t*(3) = -1.38, p =0.26; magenta squares, pre intercept awayIF = 781.4 ms and post intercept awayIF = 798.4 ms, *t*(3) = -1.35, p = 0.27).

Fig. 3a shows decision accuracy collapsed across coherences for toIF (cyan) and awayIF (magenta) sides for monkey S (triangles) and monkey B (circles) before and after 23 unilateral SC inactivation experiments. For trials with evidence favoring toIF decisions, accuracy dropped from 83% to 64% for monkey B (*t*(10) = 5.599, p = 2.28 × 10^−4^) and 79% to 59% for monkey S (*w*(11) = 0, p = 0.002). For trials with evidence favoring awayIF decisions, accuracy increased from 79% to 90% for both monkey B (*w*(10) = 66, p = 0.003), and monkey S (*t*(11) = -7.967, p = 7.00 × 10^−6^). Unilateral SC inactivation however, impairs visual and attentional processing and saccade generation^3,7,8^. Therefore, one possibility is that the biased decision-making stems from an impairment in visual or attentional processing of the choice target location or generating the movement and not decision-making processes *per se*. We think however, that these interpretations are unlikely, as on average, muscimol did not impact the slope of the psychometric function which is associated with perceptual sensitivity, and both monkeys reported toIF decisions in the high coherence conditions indicating that they could see the choice targets and make those saccades. The change in performance in the decision task suggests instead that SC inactivation impairs the balance of evidence for a decision rather than visual or attentional processing or the ability to report the decision. In the case of high coherence, the relative levels of activity between the two SCs remains greater for toIF decisions after muscimol compared to the lower coherence trials^9,10^. Nevertheless, we tested the attention and motor impairment hypotheses directly below.

**Fig. 3.**
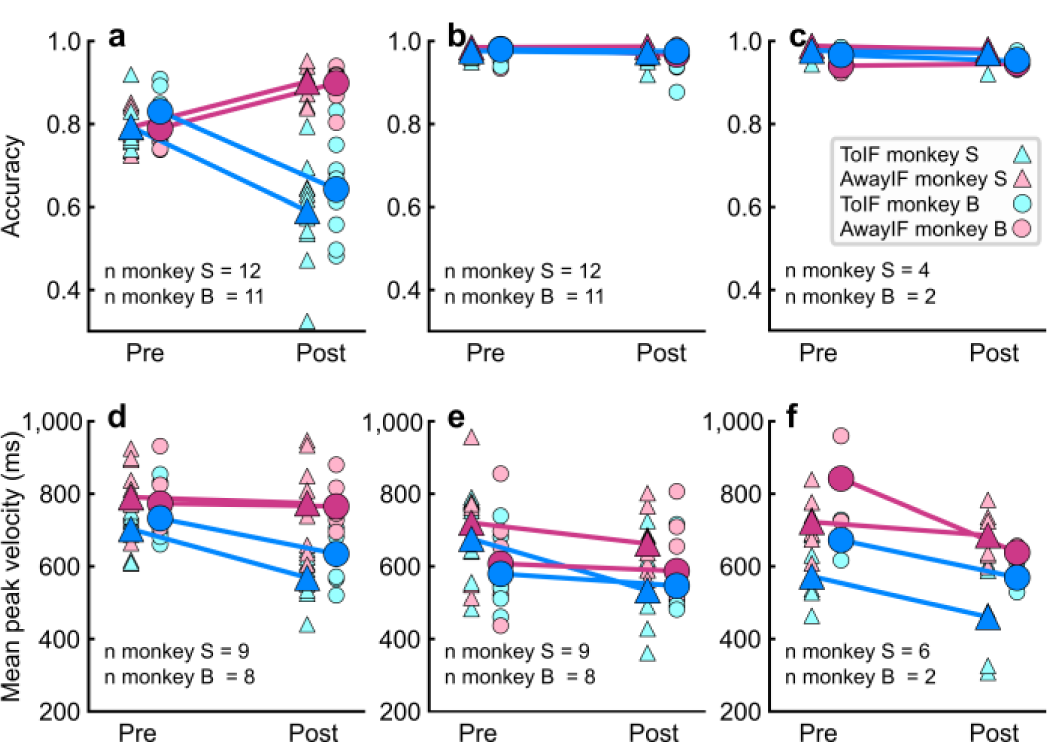
Decision but not selection accuracy is altered after SC inactivation. **a** Proportion correct (accuracy) in the delay and RT versions of the decision task is plotted for toIF trials (cyan) and awayIF trials (magenta) for 23 muscimol experiments in two monkeys, collapsed over coherence, before and after inactivation. Dark filled symbols show the mean accuracy from all experiments and less saturated colors show means of individual experiments. **b** Accuracy in the saccade selection task for the same experiments and monkeys. **c** Same as in b for the 6 saline injections in the two monkeys. **d** Mean peak saccadic velocity for the decision task before and after muscimol for toIF and awayIF decisions for 17 injections in two monkeys. Six datasets were excluded due to technical issues with the eye tracker that impacted measurement of eye speed but not assessment of choice or RT. **e** Same as in d for the saccade selection task. **f** Same as in d and e for the visually-guided saccade task used to measure saccadic velocity. There are fewer points in this plot because there were fewer saccades in this task that had the same vector target and saccade as in the decision and selection tasks. Note we also did not perform statistics with these data because of the fewer points. The data are useful for visual comparison. All statistics for accuracy and saccadic velocity appear in Extended Data Table 3.

## Decision but not selection accuracy is altered after SC inactivation

To rule out an interpretation based on attentional or motor bias or impairment, monkeys performed a selection task in which they prepared and made saccades to the same two target locations as in the decision task, but without the need for the transformation of an orientation decision to a saccade (Fig. 1c). Performance accuracy in the selection task did not change significantly after muscimol injection for either monkey for all 23 injections^5,11^ (Fig. 3b). Both monkey S and monkey B’s accuracy for toIF decisions changed from 99% to 98% after muscimol (monkey S, *w*(11) = 28, p = 0.656; monkey B, *w*(10) = 5, p = 0.249). Unilateral saline injections also produced statistically indistinguishable changes in accuracy in the selection task for toIF decisions (Fig. 3c; monkey B, 97% to 95%, *w*(10) = 1, p = 0.655; monkey S, 99% to 98%, *w*(11) = 0, p = 0.317).

Since the decision task is more difficult than the selection task, it is possible that monkeys opt to make the unaffected, awayIF saccade in the decision task more often than in the selection task. If true, we reasoned that we would see no bias in the decision task at the easiest toIF coherences, such as the 36% and 50% coherences that were near 100% accuracy pre-muscimol, since there would be no need to opt for the easier saccade if the trial was not difficult. Yet, we still observed a pronounced change in accuracy post-muscimol even for the 36% coherence condition, a finding that is better explained by a change in aspects of decision-making, rather than a motor preference under uncertainty. Also, if we assumed the monkeys opted for easier awayIF saccades, we reasoned that toIF saccades in the decision task should be harder to make and therefore slower, than saccades in the selection task, given that slower saccades indicate reduced vigor^12^. Interestingly, in some cases, the velocity of toIF saccades in the decision task was *higher* than those in the selection task after muscimol, despite matched metrics (cf., cyan symbols in post in Fig. 3d and e). The mean peak toIF saccadic velocity post-muscimol was 563.54°/s for monkey S and 634.58°/s for monkey B in the decision task. In the selection task, the mean peak toIF saccadic velocity post-muscimol was 529.00°/s for monkey S and 546.78°/s for monkey B. The decision task had significantly greater saccadic velocities than those in the selection task in six out of eight muscimol injections in monkey B, and four out of nine for monkey S (Extended Data Table 3). Fig. 3f shows a subset of the data in which monkeys made visually-guided saccades in the task used to map changes in saccade velocities. Simple, visually-guided toIF saccades also tended to be slower than those measured in the decision task, although we did not perform statistics as the data from the simple saccade task are fewer. Taken together, the higher saccadic velocities in the decision task compared to the selection task, the profound change in accuracy in the decision task, and the lack of change in accuracy in the selection task, support the embodied cognition model of decision-making and show that decision biases in the Glass pattern decision task do not stem from simple motor or attentional biases.

## SC inactivation alters evidence accumulation

We simulated multiple DDM variants with only specific parameters varying, to illustrate predicted changes in decisions and mean RTs that may occur with alterations in different aspects of decision-making (Fig. 4, Extended Data Fig. 4, Supplementary Methods). Three possibilities can be ruled out based on visual comparison of model predictions and observed data. First, the data from both monkeys are inconsistent with a symmetric, multiplicative decrease in drift rate as the slope of the psychometric function showed little to no change with SC inactivation (cf., simulation in Fig. 4a-d, simulation in Extended Data Fig. 4a-d and actual data Fig. 4q-x). Second, a symmetric increase in the toIF and awayIF boundaries is also unlikely to explain the effect of muscimol since the predicted slight changes in sensitivity and the predicted increases in correct and error RT’s for both toIF and awayIF decisions, do not match the observed results (cf., simulation in Extended Data Fig. 4q-t and actual data in Fig. 4q-x). Third, we can rule out a model based only on non-decision time, as this predicts no change in the psychometric function and joint changes in the mean RT, neither of which were observed in the post-muscimol (cf., simulation in Extended Data Fig. 4u-x and actual data Fig. 4q-x). The simulations with a decrease in the relative evidence for toIF decisions (the mean drift rate across coherences; simulations in Fig. 4m-p and Extended Data Fig. 4m-p), proportional start point change (simulations in Fig. 4e-h and Extended Data Fig. 4e-h), and an increase in the toIF boundary (simulations in Fig. 4i-l and Extended Data Fig. 4i-l), are the only parameter changes by themselves that can explain the shift in the psychometric functions that we observed in the data after muscimol (Fig. 4r,v). However, a proportional start point change does not explain the observed increases in mean error RTs and predicts a large decrease in mean correct RTs for toIF decisions which we did not observe (cf., simulations in Fig. 4g,h and Extended Data Fig. 4g,h and actual data in Fig. 4s,t,w,x). An increase in the toIF boundary may explain the changes observed in the mean correct RTs (cf., simulations in Fig. 4k,l, and Extended Data Fig. 4k,l and actual data in Fig. 4s,t,w,x), but fails to explain the magnitude of the rightward shift in the psychometric function we observed in the post-muscimol data (cf., simulations in Fig. 4j and Extended Data Fig. 4j and actual data in Fig. 4r,v). Therefore, simulations show qualitatively that a change in the mean drift rate across coherences favoring awayIF decisions, explains most of the observed data.

**Fig. 4.**
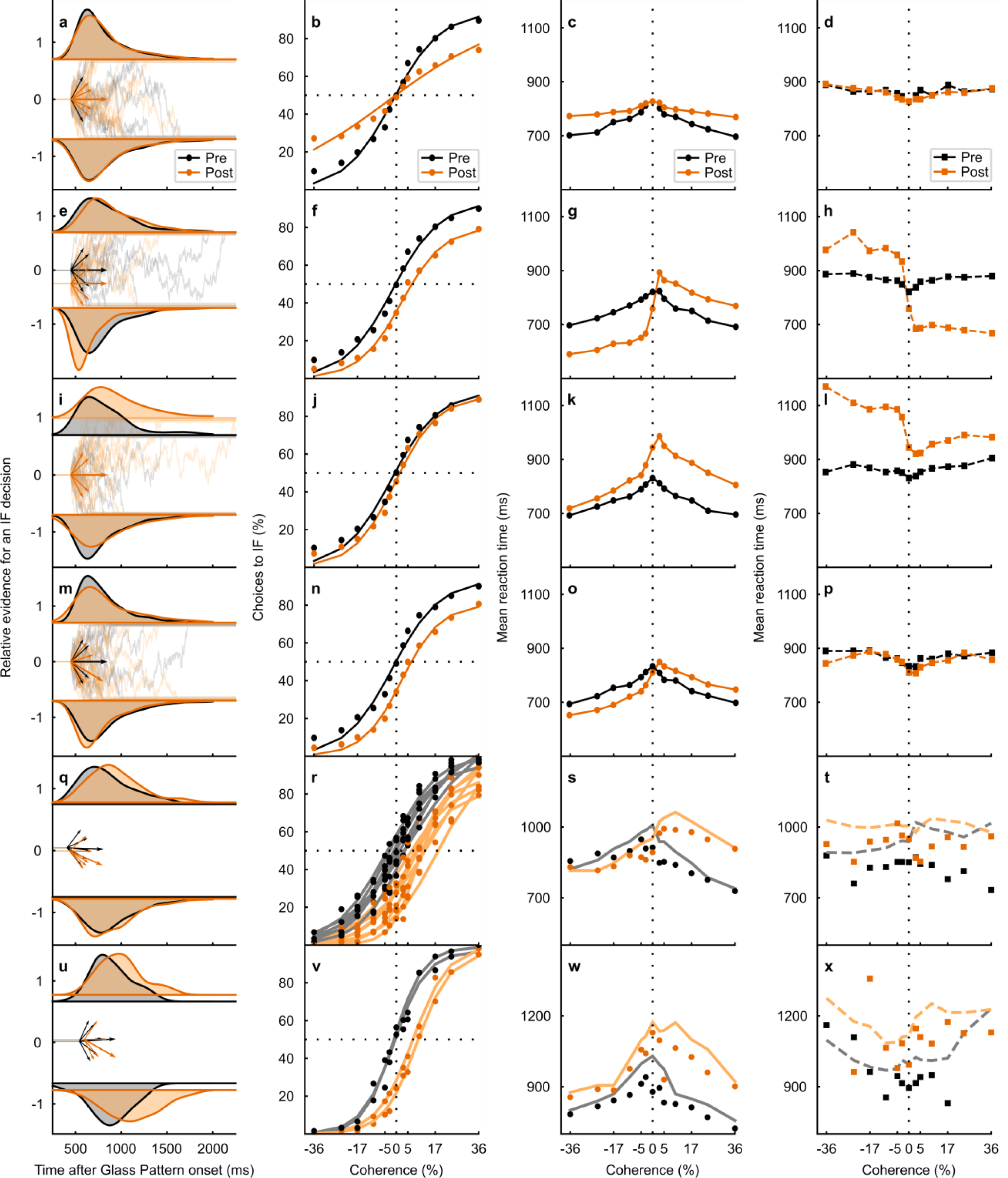
Unilateral SC inactivation alters the mean drift rate across coherences. **a** RT distribution (black = pre- and orange = post-muscimol) from the 0% coherence condition (density approximated through kernel smoothing) predicted by a DDM simulation with only a symmetric decrease in the gain of the post-muscimol drift rates. Below the RT distributions, the relative evidence for toIF decisions is plotted over time from Glass pattern onset and the short arrows show drift rate paths for toIF decisions (positive) and awayIF decisions (negative) pre- and post-muscimol, for the 0%, 10%, and 36% coherence conditions. The longer arrow shows the mean drift rate across coherences (the mean drift rate over all toIF and awayIF coherences^15^). **b** The psychometric function, plotted as a proportion of toIF decisions over coherences, predicted by the model simulation with a symmetric decrease in the gain of the post-muscimol drift rates. **c** Mean RT predictions for correct trials for each coherence condition for the DDM variant with a symmetric decrease in the gain of the post-muscimol drift rates. **d** Same as in c but for error trials. **e-h** Same as in a-d but for the DDM variant with only a change in proportional starting point of the evidence accumulation away from the IF. **i-l** Same as in a-d but for the DDM variant with an increase in the upper boundary but no absolute start point change. **m-p** Same as in a-d but for the DDM with only the same additive shift in all drift rates away from the IF, representing a change in the mean drift rate across coherences. **q** RT distributions from the 0% coherence condition of the actual data from monkey S, along with the fitted HDDM parameters of the drift rates and the mean drift rate across coherences below. Before muscimol, the mean drift rate across coherences was not significantly different than 0 (BF = 0.08) whereas after muscimol it was biased away from the IF (BF = 3.19 × 10^6^). **r** Psychometric function of the RT task sessions (n = 7) in monkey S. The circles show the data and the lines show the HDDM fits to the data. The change in the mean drift rate across coherences is evident as a rightward shift in the psychometric function post-muscimol (orange). **s** The mean RT for correct trials (circles) for all RT task data from monkey S are plotted against coherence along with their HDDM fits (lines). **t** Same as in s but for mean RT for error trials (squares) and their HDDM fits (lines). **u-x** Same as in q-t for monkey B showing all RT task data (n = 2) and the HDDM fits. Monkey B’s data and HDDM fit also show that the mean drift rate across coherence was not significantly different than 0 pre-muscimol (BF = 0.14) but significantly differed from 0, away from the IF, post-muscimol (BF = 17.87).

We next fit hierarchal and non-hierarchal drift-diffusion model variants (HDDM and DDM) and urgency-gating model varients^13,14^ (UGM) to the performance and RT data (seven in monkey S and two in monkey B; Methods and Supplemental Methods). Parameter estimation of pre- and post-muscimol data indicated that multiple parameters changed after muscimol (Extended Data Fig. 5). However, the only consistent parameter change in both monkeys across experimental sessions that explained the rightward shift in psychometric functions was an increase in the relative awayIF evidence accumulation as reflected by the mean drift rate parameter (Fig. 4q,u arrows, Extended Data Fig. 5). The mean drift rate across coherences differed from 0 after muscimol for both monkeys (HDDM, monkey S, Bayes factor (BF) = 3.19 × 10^6^; monkey B BF = 17.87) but not before muscimol (monkey S BF =0.08; monkey B BF = 0.14). The posterior probability of a negative change in mean drift rate across coherences was 99.7% in monkey S (posterior medians pre = -0.06, post = -0.64) and 99.0% in monkey B (posterior medians pre = 0.09, post = -0.85). The start point of evidence accumulation did not differ from 0.5 after muscimol in either monkey (HDDM, monkey S, BF=0.09; monkey B, BF=0.35), nor before muscimol (monkey S, BF = 0.73; monkey B, BF = 0.18). The posterior probability of a proportional start point change away from the IF, was 95.1% in monkey S (posterior medians pre = 0.54, post = 0.49; probability of a toIF start point change was 100% - 95.1% = 4.9%), but was 70.6% for a proportional start point change toward the IF in monkey B (posterior medians pre = 0.52, post = 0.55; probability of an awayIF start point change was 100% - 70.6% = 29.4%), indicating opposite starting point changes in both monkeys. The posterior probability of a non-decision time increase was 94.5% in monkey S (posterior medians pre = 408 ms, post = 433 ms) and 97.0% in monkey B (posterior medians pre = 543 ms, post = 597 ms). The posterior probability of a symmetric boundary increase was 78.9% in monkey S (posterior medians pre = 1.5, post = 1.6) and 95.2% in monkey B (posterior medians pre = 1.3, post = 1.5). We found little evidence for a change in the proportion of lapse trials, trials in which decisions are determined randomly, for either monkey (a positive change in lapse proportion was 72.5% in monkey S with posterior medians pre = 0.32 and post = 0.38 and 54.0% in monkey B with posterior medians pre = 0.45 and post = 0.46). The non-hierarchical DDM fits also showed the same patterns of parameter changes indicating that the results are robust to modeling methods (Extended Data Fig. 5k,l). Overall, the mean drift rate across coherences was the only parameter that changed significantly (> 95% posterior probability) after SC inactivation in both monkeys. Likely (> 94.5%) non-decision and somewhat likely (> 78.9%) symmetric boundary increases were observed in both monkeys, but neither parameter explains the rightward shift in psychometric functions we observed in the post-muscimol data.

To test directly which parameter change best explained the effect of SC inactivation on decision-making, we fit HDDM variants with the following parameters free to vary while keeping all others fixed to the observed data: mean drift rate across coherences (HDDM-*Δ*), proportional start point (HDDM-*w*), non-decision time (HDDM-*τ*), and proportional start point along with bound (HDDM-*a,w*, to test the fitting of either single or symmetric bound changes; Supplementary Methods). An HDDM fit with only the mean drift rate across coherences allowed to change across pre- to post-muscimol explained the shift in psychometric function almost as well as the HDDM fit with all parameters allowed to change (HDDM-*Δ* explained 97.6% of the variance of the psychometric function and the full HDDM explained 98.3% for monkey S; the HDDM-*Δ* explained 98.3% of the variance and the full HDDM explained 99.3% for monkey B) and fit the shifts in psychometric function better than all other competing models (Extended Data Fig. 6, Extended Data Table 4).

We also fit an urgency-gating model^13,14^ (UGM) to the data for both monkeys to assess whether our findings were robust to different decision-making model assumptions and to see whether an urgency signal change might also explain the effect of SC inactivation (Supplementary methods). Fig. 4q,u and Extended Data Fig. 6 also show that the pre-muscimol RT distribution of monkey B has a more symmetric shape, relative to monkey S, which the UGM predicts. Although evidence for the best fitting model type was mixed across monkeys (Extended Data Table 4), only a decrease in the mean drift rate across coherences and the urgency slope parameter decrease were consistent in both monkeys (Extended Data Fig. 5k-l). However, allowing drift rates to vary from pre- to post-muscimol (Supplementary Methods) explained more of the in-sample variance in the psychometric function of both monkeys (UGM*-δ* explained 87.8% for monkey S and 83.1% for monkey B, Extended Data Table 4) than UGMs with urgency slope allowed to vary, a parameter that also influences response proportion and RT (UGM-*m* explained 20.0% for monkey S and 26.2% for monkey B, Extended Data Table 4). Taken together, in both the DDM and the UGM, a change in the mean drift rate across coherences best explains the effect of unilateral SC inactivation on perceptual decision-making.

Our results are based on assuming a one dimensional accumulator model. However, an alternative possibility is that there are two independent accumulators^25^ and SC inactivation impairs evidence accumulation in one, perhaps by altering its gain^24^. In the two dimensional model, the gain of the toIF accumulator would decrease with unilateral SC inactivation and the gain of the awayIF accumulator would remain unchanged. To test the two independent accumulator with gain model, we fit the full HDDM with the same parameters but with drift rates constrained to a linear comparison of two accumulators (Supplementary Methods). We found strong evidence for a gain decrease on the toIF accumulator post-injection. The BF of *G*_*(toIF)*_ not equal to one was estimated to be very large (>10^307^) with a posterior median of *G*_*(toIF)*_ = 0.6217. We found no evidence for gain decreases in any other experimental conditions; recovery, pre-saline, post-saline, BF of *G*_*(toIF)*_ not equal to one ranged between 0.0114 to 0.2232 and the posterior medians of *G*_*(toIF)*_ ranged from 0.9307 to 1.0832. These results provide evidence that inactivation of the SC impairs evidence accumulation for perceptual decisions by altering the gain of evidence accumulation in one of two competing accumulators.

## Conclusion

We provide compelling evidence that unilateral inactivation of the primate SC alters perceptual decision-making, not by changing sensory or motor processing, nor by biasing evidence before accumulation begins. Rather, unilateral SC inactivation shifts the balance of evidence accumulation away from the IF. This is similar to changing the decision criterion in the SDT framework consistent with results in rodents and our previous results in monkeys^16,17^. Current conceptions of perceptual decision-making fall into two main categories; one in which sensory evidence is evaluated, categorized and then forwarded to motor areas to guide choices of action^18^, and a second in which brain areas involved in getting ready to act are the same areas that accumulate evidence over time to form a decision; referred to as embodied cognition. Our results support an embodied cognition model of perceptual decision-making. The SC is well-known for its role saccade preparation and generation^7,19^, and we show here that toIF saccades remain relatively intact with SC inactivation^5,11^, in spite of significant alterations in decision-making about the orientation of a Glass pattern.

Our results suggest exciting new possibilities for how perceptual decisions are formed and converted to choices of action in the brain. In primates, including humans, and in rodents, perceptual decisions are thought to arise from evidence accumulation in forebrain areas such as area LIP (PPC) and dlPFC (FOF). Our results indicate that the SC plays a causal role in how evidence is accumulated - wherever it is accumulated. A very real possibility is that each SC contains an independent accumulator^25^ and inactivation of one SC, as we performed here, shifts the balance of evidence toward the other accumulation process and decision. Furthermore, as suggested by our two-dimensional modeling results, inhibition of the SC behaves as a gain element that controls evidence accumulation that occurs either in the SC or elsewhere. Recent evidence from experiments inactivating the forebrain accumulators, LIP (PPC) and dLPFC (FOF) however, call into question the causal role of these areas in evidence accumulation^18,20-23^ and suggest that FOF in rodents at least, participates in decision-making after evidence has been accumulated^22^. Therefore, we propose that critical evidence accumulation and transformation of a decision to a choice of action happens in the SC, close to the motor output and that the SC also impacts evidence accumulation occurring elsewhere in the brain, perhaps through feedback circuits via thalamic nuclei from the SC^7^. This exciting possibility awaits further investigation.

## METHODS

### Surgery

We implanted two adult male rhesus monkeys (Macaca mulatta), weighing between 9-11 Kg (monkey S 15yo and monkey B 11yo) with eye loops for measuring eye position^24^, a post for stabilizing the head and a recording chamber for accessing the superior colliculus (SC). Devices were placed using MRI-guided surgical software (Brainsight, Rogue Research, Montreal, CA) and stereotaxic coordinates (0ML, -3AP, angled 38° posteriorly). All surgical procedures were performed under general anesthesia using aseptic procedures and all surgical and experimental procedures were approved by the UCLA Chancellor’s animal research committee and complied with and generally exceeded standards set by the Public Health Service policy on the humane care and use of laboratory animals as well as the American Primate Veterinarian Guidelines.

### Behavior and electrophysiology

We used a real-time experimental control and visual stimulus generation system, REX and VEX, developed and distributed by the Laboratory of Sensorimotor Research National Eye Institute (Bethesda, MD) to create the behavioral paradigms^25^. We used the magnetic induction technique^26^ (Riverbend instruments, Birmingham, AL) and the EyeLink 1000 eye tracker system (SR Research Ontario, CA) to measure voltage signals proportional to horizontal and vertical components of eye position (monocular mode; 2kHz). Eye position signals were low-pass filtered (8 pole Bessel - 3dB, 180 Hz; Bak Electronics; Umatilla, FL) and digitized at 16-bit resolution and sampled and saved to disk at 30 kHz using Blackrock Microsystems NSP hardware system controlled by the Cerebus software suite (Blackrock Microsystems, Salt Lake, UT). We used an automated procedure to define the onset of saccadic eye movements using eye velocity (20°/s) and acceleration criteria (5000°/s^2^). The adequacy of the algorithm was verified and adjusted as necessary on a trial-by-trial basis by the experimenter. We omitted < 10% of trials from one monkey because of saccadic festination (i.e., small saccades toward the target followed by larger ones).

Two trained monkeys performed three behavioral tasks: 1) a visually-guided saccade task to map response fields (RF) before muscimol injections as well as obtain changes in saccade velocity after muscimol, 2) a selection task to measure saccadic motor preparation and bias before and after muscimol injection, and 3) a task for assessment of perceptual decision-making performance before and after muscimol. Pre- and post-muscimol data were collected on the same day and recovery data were collected ∼24 hours after muscimol injection. Recovery data appear in Extended Data Figs.1 and 3 and the associated statistics in Extended Data Table 2.

### Response field (RF) mapping

We used a visually-guided saccade task to map RFs of SC sites. A red fixation spot appeared (14.32 cd/m^2^ for monkey B and 22.37 cd/m^2^ for monkey S) and monkeys maintained fixation at this location for a random time of 500-1000 ms with an accuracy of 3.5° square determined by an electronic window. Next, a white spot (48.27 cd/m^2^ for monkey B and 109.76 cd/m^2^ for monkey S) appeared in the periphery. Monkeys remained fixating centrally for a random time of 800-1200 ms until the fixation spot disappeared, cueing the monkeys to look at the spot in the periphery. If monkeys looked at the peripherally-located spot within a 4.5° square determined by an electronic window, they received a sip of water or preferred juice for reward. Incorrect saccades were not rewarded. The saccade target was positioned manually and pseudorandomly throughout the visual field (Fig. 1d).

While monkeys performed the visually-guided saccade task, we recorded single and multiple neurons in the intermediate layers of the SC using custom injectrodes that allowed for neuronal recording and injection of compounds simultaneously (Fig. 1e). Injectrodes were inserted through a guide tube positioned by a grid system^27^ and were moved in depth by an electronic microdrive system controlled by a graphical user interface on a PC running Windows (*Nan Instruments*, Israel). Action potential waveforms were bandpass filtered (250 Hz to 5 kHz; 6 pole Butterworth) and amplified by a differential amplifier and then sampled, digitized and saved to disk at 30 kHz with 16 bit resolution using the Blackrock NSP hardware system controlled by the Cerebus software suite for offline sorting as necessary (Blackrock Microsystems, *Salt Lake, UT*). When possible, neurons were isolated online using time and amplitude windowing criteria and the times of action potentials were saved to disk similarly. Response fields (RF) of SC neurons (either single neurons, if well-isolated, or multiple neurons, if not) were mapped during the experiment using customized MATLAB scripts (MathWorks *Natick, MA*) that plotted the average discharge rate from 50ms before to 50ms after the saccade onset for each target position (Fig. 1e). We considered the center of the RF to be the location at which a saccade was associated with maximal discharge (audibly, visually and quantitatively). Only locations with RF eccentricities greater than 8° were included to ensure no overlap of the RF with the center of the visual field.

A variant of this task allowed us to measure peak saccadic velocity before and after muscimol injections. This task had randomized delay times taken from a truncated exponential distribution (fixation time mean 400 ms, range 320-560 ms; delay-time mean 800 ms, range 640-1120 ms) and fixed target positions (Fig. 1f).

### Selection task

The second task was a visually-guided, delayed-saccade task in which two isoluminant targets (14.65 cd/m^2^ for monkey B or 21.38 cd/m^2^ for monkey S) appeared in the periphery. One target was located at the center of the RF and the other was located 90° opposite in the other hemifield (Fig. 1c). One target was red and the other was white. The position of the red and white targets switched randomly on each trial. After the fixation point (48.62 cd/m^2^ for monkey B or 106.86 cd/m^2^ for monkey S) appeared, monkeys remained fixating on this spot for a mean delay time of 400 ms (320-560 ms, truncated exponential) until the targets appeared. A second mean delay of 800 ms (640-1120 ms, truncated exponential) occurred and then the fixation spot disappeared, cueing the monkey to look at the white target. If the monkey looked at the white target with an accuracy of 5.5° determined by an electronic window, it received a sip of preferred juice or water for reward. For 12 of the 23 muscimol experiments and 2 of 6 saline experiments we used fixation delays of 100 ms and a mean delay-period of 350 ms (200-500 ms, truncated exponential). For the latter data, we analyzed only those trials with a 400 ms or greater delay-period. This task required the same attentional allocation to the target location as well as the same motor preparation as the decision task, however, it did not require the transformation of the Glass pattern orientation to the saccade location as did the decision task, allowing us to assess impairments in motor preparation and biases in saccades to the left or right hemifields before and after the muscimol injection.

### Glass pattern decision-making task

To assess perceptual decision-making performance before and after muscimol injections, monkeys performed a one interval, two-choice, perceptual decision-making task in which they reported the orientation of a dynamic Glass pattern (decision task)^1^. Monkeys reported their decisions by making saccades to a target located in the left or right hemifield corresponding to the orientation of the perceived Glass pattern. We parameterized the difficulty of the decision by varying the coherence of the Glass pattern among 2 sets of coherences: 0%, 5%, 10%, 17%, 24%, 36%, 50% performed by monkey S for 3 muscimol experiments on the delay decision task, and 0%, 3%, 5%, 10%, 17%, 24%, 36% performed by both monkeys for all other experiments, including both muscimol and saline and delay and RT task versions. Monkeys received water or preferred juice reward for correct trials and on the 0% coherence trials, they received reward on half of the trials randomly. Once monkeys were well-trained and after performing the first three muscimol experiments on a fixed ratio (FR) 1 reward schedule (rewarded on every correct trial), monkeys performed the task on a variable ratio (VR) schedule such that on average, but with some variation, every 3^rd^ or 5^th^ correct trial received reward to encourage consistent performance (VR3 or VR5)^28^.

Trained monkeys performed the decision task in a delayed version and a reaction time (RT) version. In the delayed version, a fixation spot appeared (2.93 cd/m^2^ for monkey B and 9.62 cd/m^2^ for monkey S) at the center of the display. After a mean 300 ms delay (240-420 ms, truncated exponential), two isoluminant choice targets appeared (3.06 cd/m^2^ for monkey B or 6.36 cd/m^2^ for monkey S). After another mean delay of 700 ms, (560-1400 ms, truncated exponential), the Glass pattern cue appeared at the location of the fixation point and remained illuminated for 950 ms (760-1900 ms, truncated exponential). After the Glass pattern cue disappeared, there was a delay-period with mean of 800 ms (640-1120ms, truncated exponential). The removal of the fixation point cued the monkeys to report their decision by looking at one of the two choice targets. Monkeys remained fixating at the correct choice target for a mean of 350 ms (280-490 ms, truncated exponential), before receiving fluid reward. Monkeys performed a variant of the delayed task with a shorter delay period (50-100 ms) for 12 muscimol experiments and two saline experiments (Fig. 1a).

The RT version of the decision task was identical to the delay version except that, in the RT task, the Glass pattern appearance and the removal of the fixation spot occurred simultaneously and the monkeys reported their decisions at any time (Fig. 1b). To discourage fast guessing, we implemented a fixed time to reward (900 ms monkey B, 1000 ms monkey S) and a RT-dependent inter-trial interval^29^. The results of muscimol injections on choice behavior were similar in both the delay and RT versions of the task, so the data are collapsed unless otherwise indicated. Modeling of RTs and choice behavior is based on data only from the RT task for each monkey separately (Fig. 4q,u).

### Statistical tests of significance

We analyzed all the data using customized scripts developed in MATLAB 2016b (MathWorks *Natick, MA*), Python 3 (3.7.6), R (v.3.4.0), and IBM SPSS Statistics 25. To assess for significant differences between the *α, β*, RT mean collapsed across coherences and injections, RT slope, and RT intercept parameters of pre- and post-injection, and pre-injection and recovery, we performed paired two-tailed *t*-tests (if both sessions data sets were normally distributed using Lilliefors or Shapiro-Wilk test) or Wilcoxon Signed rank tests (if either of the sessions were non-normally distribution) for pre- vs. post-injection and pre-injection vs. recovery comparisons with Bonferroni corrections (2 comparisons, *α* = 0.05/2 = 0.025), where the pre- vs post-injection comparisons appear in the main text and the pre-injection vs recovery comparisons appear in Extended Data Table 2. The same tests were performed for the analysis of accuracy data from the decision and the selection tasks for toIF and awayIF, except that with the Bonferroni corrections for 4 pairwise comparisons tests (Extended Data Table 3), significance cut-off value *α* is 0.05/4 = 0.0125. For analyzing the post-muscimol peak saccadic velocity for toIF saccades between the decision task and the selection task, we performed unpaired *t*-tests (for normal data, two-tailed) or Wilcoxon Rank Sum (if non-normally distributed) with Bonferroni correction of *α* = 0.05/9 = 0.0056 for the tests for monkey S, since 9 t-tests were performed on data from each injection, and *α* = 0.05/8 = 0.0063 for the tests for monkey B, since 8 t-tests were performed on data from each injection, with a total of 17 muscimol injections between the 2 monkeys (Extended Data Table 3). Six injections were excluded due to technical issues with the eye tracker that impacted measurement of eye speed but not assessment of choice or RT.

### Modeling and simulations

We fit Signal Detection Theory (SDT), drift-diffusion (DDM) and urgency-gating models (UGM) to the behavioral data to understand how decision-making was impacted by unilateral inactivation of the SC^13,14,30,31^. SDT is a static model of decision-making and makes predictions only about choice performance, whereas the DDM and UGM are dynamic models that make predictions about both choice performance and RTs. Parameters of decision-making models are thought to be instantiated by aspects of neuronal activity and many have been verified in human behavioral experiments^32,33^. We performed simulations, model fitting and model comparisons using customized scripts developed in MATLAB 2016b (MathWorks *Natick, MA*), Python 3 (3.7.6), JAGS^34^ (4.3.0), and R (v.3.4.0). We also used the published libraries Palamedes^35^, pyjags^36^ (1.2.2), the Wiener module for JAGS^37^, and ChaRTr^38^. For equations, simulations, parameter estimation, and model comparison results, see Supplementary Methods, Extended Data Fig 4-6, and Extended Data Table 4.

### Parameter estimates in SDT

We fit a two parameter logistic function using the Palamedes toolbox to the choice performance data for each monkey for all experiments using the equation:

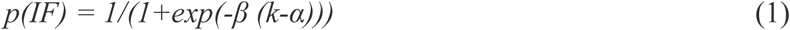

where *p(IF)* denotes the proportion of choices to the inactivated field (IF) for each coherence condition (*k*). *α* and *β* are free parameters determined using maximum likelihood methods and provide measures of decision bias and sensitivity of the psychometric function respectively^35,39,40^. Fig. 2 and Extended Data Fig. 3 show the two parameter model fits and results. Extended Data Fig. 2 shows comparisons between the two, three and four parameter model fits. Because there were no differences in the quality of the fits for the three models, we opted to use the simper fewer parameter model for the analysis. For three of the muscimol injections, we used 50%, 36%, 24%, 17%, 10%, 5%, and 0% coherence conditions for the fits. For the other 20 muscimol sessions and 6 saline injection sessions, we used 36%, 24%, 17%, 10%, 5%, 3%, and 0% coherence conditions. We also calculated *d’* and *c*, using SDT equations as described in Macmillan and Creelman^30^ and Supplementary methods (S1-2) and these results appear in Extended data Fig. 1.

### Parameter estimates in drift-diffusion Models (DDMs)

We fit a hierarchical DDM (HDDM) and a non-hierarchical DDM (DDM) that describe RT and choice distributions. Non-decision time (*τ*) is the sum of visual and motor processing. The decision boundary (*a*) determines the amount of evidence needed to make a decision. The starting point of evidence accumulation (*w*) was fit as a proportion of the boundary parameter and describes the initial bias in evidence accumulation before a trial begins, where *w* = 0.5 indicates the start point is the middle between the upper (toIF) and lower (awayIF) bound, *w* > 0.5 indicates the start point is closer to the upper (toIF) bound, the *w* < 0.5 indicates the start point is closer to the lower (awayIF) bound. The drift rate (*δ*) is the average evidence accumulation rate during a trial and is driven by the strength of the evidence extracted from the Glass pattern stimulus. There is also a mean drift rate across coherences and directions (Δ), also known as the “drift-criterion” or the “zero point of drift rate^31^”. The lapse rate (*λ*), was fit and defined as the percentage of trials in which choices were determined randomly. The lapse rate was only fit in the HDDM but not the DDM.

We used both hierarchical Bayesian methods (HDDM with *k* = 13 coherence conditions) and Quantile Maximum Probability Estimation (QMPE; DDM with *k* = 11) to estimate parameters. Both estimation methods yielded similar parameters (Extended Data Fig 5). Hierarchical Bayesian methods were applied to fit the HDDM to the data from the RT task from both monkeys: 7 muscimol and four saline sessions from monkey S and 2 muscimol sessions from monkey B. We used JAGS to draw samples from posterior distributions using Markov Chain Monte Carlo (MCMC) samplers^34^. Hierarchical mean parameters per monkey and experimental condition (pre-, post-, and recovery for muscimol and saline) were assumed to better fit the data from each experimental session (Supplemental methods equations S9-15 and Extended Data Table 4). We also fit HDDM with only drift rate, proportional start point, non-decision time, or proportional start point with the bound varying across conditions (equations S16-44 in Supplementary Methods). Parameter estimates for full hierarchical models appear in Extended Data Fig. 5a-j.

To calculate the probability of change in mean drift rate across coherences, we estimated the posterior distributions using kernel density estimation and then summed the density from the lowest negative sample to zero. We also calculated Bayes Factors (BF) using the Savage-Dickey density ratio. BF describes the relative evidence of the mean drift rate (*Δ*) not equal to zero. The BF for mean drift rate was the ratio of the prior density at *Δ* = 0 over the posterior density at *Δ* = 0. We also calculated the BF for whether the proportional start point was different than 50% of the relative evidence units required to make a decision (the boundary was also a free parameter). The BF for hierarchical initial bias was calculated as the ratio of the prior density at *w* = 0.5 over the posterior density at *w* = 0.5. In the model HDDM-*G* (supplementary methods) the BF for a gain change on one accumulator resulting from inactivation of one SC, not equal to one was calculated as the ratio of the prior density at *G*_*(toIF)*_ = 1 over the posterior density at *G*_*(toIF)*_ = 1. A BF over three is considered positive evidence for an effect (i.e. the effect is three times more likely under the alternative hypothesis than the null hypothesis) whereas, over 20 is strong evidence.

### Parameter estimates in urgency-gating model (UGM)

UGM are another class of decision-making model in which the sensory evidence is low-pass filtered to prioritize more recent evidence and then multiplied by a linearly growing urgency signal^13,14^. We used QMPE (*k* = 11) to estimate UGM parameters from the data from the RT task from both monkeys; seven muscimol sessions from monkey S and two muscimol sessions from monkey B (Supplementary Methods). We fit the UGM to pre- and post-muscimol data separately to discover which parameters changed. We also fit UGMs with an urgency slope parameter or a drift rate parameter free to vary in post-muscimol data while keeping other parameters fixed to their parameter estimates from the pre-muscimol data (Supplementary Methods). Parameter estimates for all models appear in Extended Data Fig. 5.

### Model comparisons

We generated posterior predictive samples for hierarchical drift-diffusion models (HDDMs) and predicted choice and RT distributions from non-hierarchical DDM and UGM parameter estimates using QMPE with in-sample and out-of-sample datasets generated from an 80% / 20% random data split in each experimental session. We used predicted choice and RT distributions, as well as Akaike and Bayesian Information Criteria (AIC/BIC) where applicable, to find models that best described the effect of unilateral inactivation of the SC (Supplementary Methods). Percentage variance of RT and choice statistics explained by prediction are given as derived from *R*^*2*^_*pred*_ (Supplementary Methods). In-sample and out-of-sample prediction results appear in Extended Data Table 4.

## Data Availability

The data reported in this manuscript are available from the corresponding author upon reasonable request.

## Code Availability

MATLAB, Python, R, and JAGS analysis code will be made available in the following repository https://gitlab.com/fuster-lab-cognitive-neuroscience/sc-inactivation-project upon publication.

## Acknowledgements

We are grateful to Dr. Joaquin Fuster for all of his support. We thank Dr. Jochen Ditterich for suggesting the gain hypothesis and Dr. Alex Huk for comments on a previous version of the manuscript. We thank Dr. Piercesare Grimaldi for help with the initial injection experiments, Mr. Marcus Lenoir and Mr. Dave Tokuda for monkey care, Mr. Alessandro Fabro for programming support and Dr. Richard Krauzlis for monkey illustrations. This work was supported by EY013692 to MAB.

## Author contributions

E.J.J., A.B, and M.A.B. designed the study. E.J.J., A.B., T.T., E.A. collected the data with M.A.B. guidance. E.J.J., A.B., and M.D.N. analyzed the data with M.A.B. guidance. E.J.J., A.B., M.D.N., and M.A.B. interpreted the results and wrote the paper.

## Additional Information

None

## Supplementary Information

is available for this paper.

## Reprints and permissions information

are available at http://www.nature.com/reprints.

## Extended Data Figures and Captions

**Extended Data Table 1.** Unilateral Muscimol Injections into the monkey SC. (Associated with Figs. 1 and 2 main text) Details of the injections in two monkeys. Each row shows an individual injection (n=29) and each column shows the specifics of one particular injection. The column labeled monkey indicates that the experiment was performed in monkey S or monkey B and the treatment column indicates whether the injection was saline or muscimol. The next column indicates the total injection volume in µl. The column labeled Conc. (µg/µl) lists the concentration of muscimol or saline and the side of injection column lists the side of the brain in which the injection was made. The final column indicates the version of the decision task used (RT or delay).

| Exp. # | Monkey | Treatment | Volume (uL) | Conc. (ug/uL) | Side of injection | Decision task type |
| --- | --- | --- | --- | --- | --- | --- |
| 1 | S | Muscimol | 0.75 | 0.5 | Left | Delay |
| 2 | S | Muscimol | 0.5 | 0.5 | Left | Delay |
| 3 | S | Muscimol | 0.5 | 0.5 | Left | Delay |
| 4 | B | Muscimol | 0.5 | 4.0 | Right | Delay |
| 5 | B | Muscimol | 0.5 | 4.0 | Right | Delay |
| 6 | B | Muscimol | 0.5 | 4.0 | Right | Delay |
| 7 | B | Muscimol | 0.5 | 4.0 | Right | Delay |
| 8 | B | Muscimol | 0.5 | 4.0 | Left | Delay |
| 9 | B | Muscimol | 0.5 | 4.0 | Left | Delay |
| 10 | S | Muscimol | 2.0 | 2.0 | Right | Delay |
| 11 | S | Muscimol | 1.0 | 2.0 | Right | Delay |
| 12 | B | Muscimol | 0.5 | 4.0 | Left | Delay |
| 13 | B | Saline | 0.5 | 9.0 | Left | Delay |
| 14 | B | Saline | 0.5 | 9.0 | Left | Delay |
| 15 | B | Muscimol | 0.5 | 4.0 | Left | Delay |
| 16 | B | Muscimol | 0.5 | 4.0 | Right | Delay |
| 17 | B | Muscimol | 0.5 | 4.0 | Right | RT |
| 18 | B | Muscimol | 0.5 | 4.0 | Right | RT |
| 19 | S | Muscimol | 1.0 | 2.0 | Left | RT |
| 20 | S | Saline | 1.0 | 9.0 | Left | RT |
| 21 | S | Saline | 1.0 | 9.0 | Left | RT |
| 22 | S | Muscimol | 1.0 | 1.5 | Left | RT |
| 23 | S | Muscimol | 1.0 | 1.5 | Left | RT |
| 24 | S | Muscimol | 1.0 | 1.5 | Left | RT |
| 25 | S | Muscimol | 1.0 | 1.5 | Left | RT |
| 26 | S | Saline | 1.0 | 9.0 | Right | RT |
| 27 | S | Muscimol | 1.0 | 1.5 | Left | RT |
| 28 | S | Saline | 1.0 | 9.0 | Right | RT |
| 29 | S | Muscimol | 1.0 | 1.5 | Right | RT |

**Extended Data Fig. 1.**
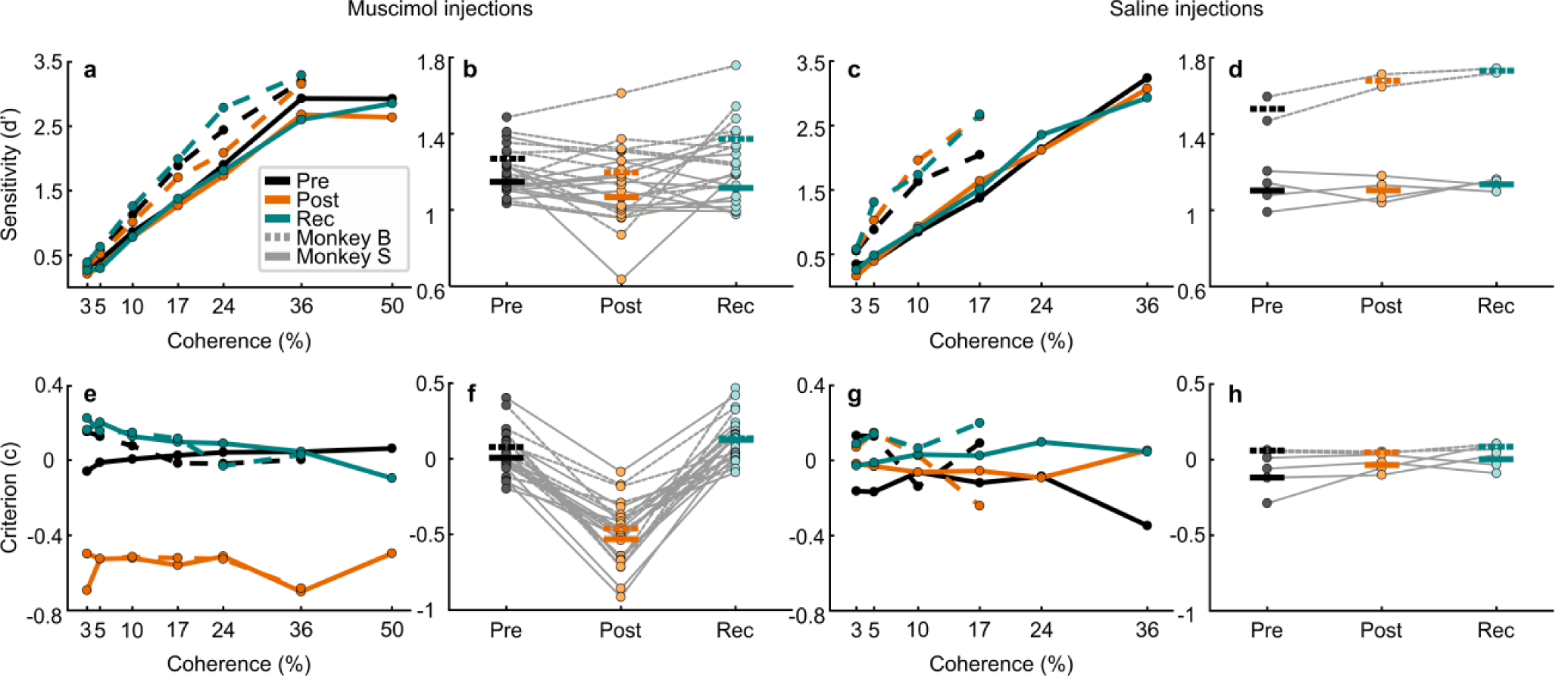
Decision criterion but not sensitivity, is impacted by unilateral SC inactivation during one-interval, two-choice perceptual decision-making. (Associated with Fig. 2 main text) **a** Sensitivity, as measured by *d’* is plotted against coherence for both monkeys pre-muscimol (black circles and lines), post-muscimol (orange circles and lines) and 24 hour recovery (green circles and lines). Dashed lines show data from monkey B and solid lines show data from monkey S. Note that for monkey S, there is an additional 50% coherence condition (Methods). Qualitatively, monkey B showed a higher sensitivity for the same Glass pattern coherences than monkey S. **b** *d’* collapsed over coherence and plotted for pre-muscimol (grey circles), post-muscimol (orange circles) and recovery (green circles) for both monkeys. Dashed lines show data from monkey B and solid lines show data from monkey S. The horizontal lines indicate the mean *d’* for each session. On average there were no significant changes in *d’* with muscimol in either monkey (monkey S, *t*(11) = 1.54, p = 0.152; monkey B, *t*(10) = 1.51, p = 0.161). **c-d** Same as in a and b for the saline injections. Because we only had two saline injections in monkey B, we collapsed the data across monkeys (n=6) for statistical analysis, but the data are shown separated by monkey. We found no significant differences in *d’* with saline (*t*(5) = -1.20, p = 0.283). Note that there are no *d’* or criterion (*c*) values for monkey B for the 24% and 36% coherences due to a lack of errors for the awayIF post-muscimol 24% and 36% coherence trials. **e** Criterion (*c*) plotted against coherence for pre-muscimol (black), post-muscimol (orange), and recovery (green). Dashed lines are from monkey B and solid lines are from monkey S. This plot is shown for symmetry with the *d’* plot although criterion changes across coherences are not particularly meaningful as monkeys are not expected to change their criterion across coherences as the coherences were randomized from trial to trial and there was no way for the monkeys to know which coherence was impending. **f** Criterion collapsed over coherence plotted for pre-muscimol, post-muscimol and recovery. For both monkeys, *c* changed significantly with muscimol (monkey S, *t*(11) = 9.34, p = 1.46 × 10 ^−6^, monkey B, *t*(10) = 7.48, p = 2.10 × 10 ^−5^). **g-h** Same as in e-f for the saline experiments. We found no significant differences in *c* with saline injections (*w*(5) = 3, p = 0.156). Consistent with the psychometric function results shown in Fig. 2, unilateral inactivation of SC with muscimol produced changes in decision bias and not perceptual sensitivity.

**Extended Data Fig. 2.**
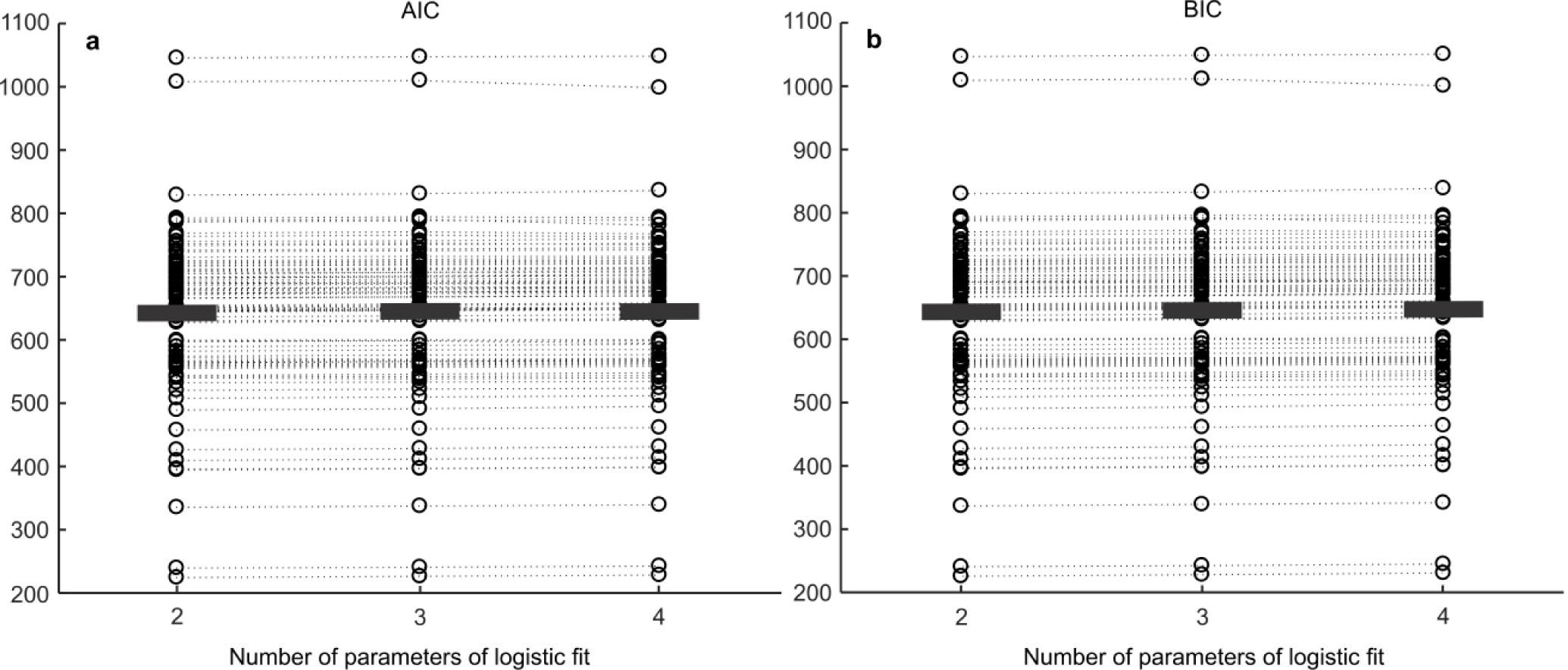
AIC and BIC scores for the two, three and four parameter logistic function fits. (Associated with Fig. 2 of the main text) **a** AIC scores for each injection for pre- and post-muscimol and recovery for all muscimol and saline injections (n = 87) for each two, three, and four parameter logistic fits to the performance data. The circles show the AIC score of the logistic fit to each individual session from the n = 87 total sessions (pre, post and recovery * 29 injections), and the black, horizontal bars show the mean AIC score. The dotted lines connect the same data sessions that were fit across the two, three, and four parameter fits to see if there were any changes in AIC score between the fits with different number of parameters. The two parameter logistic model has two parameters: *α* (decision bias) and *β* (sensitivity) following the equation *p(IF) = 1/(1+exp(-β (k-α)))* (Eq 1 in methods), which was used to fit the psychometric functions in Fig 2 and Extended Data Fig. 3. The three parameter logistic model includes: *α, β*, and *λ* (lapse rate or the difference between perfect performance and the top and bottom asymptotes) following the equation *p(IF) = γ + (1-2γ))/(1+exp(-β (k-α)))*. The four parameter logistic model includes: *α, β* and *λ* (lapse rate or the difference in perfect performance and asymptotic performance for toIF decisions) and *γ* (lapse rate or the difference in perfect performance and asymptotic performance for awayIF decisions) following the equation *p(IF) = γ + (1-γ-λ))/(1+exp(-β (k-α)))*. When looking at the AIC scores for the two, three, and four parameter fits (lower AIC scores indicate a better fit given model complexity), we see that the data are explained equally well or better with the models without lapse rates, with mean scores of 638.80 for the two parameter fit, 640.51 for the three parameter fit, and 641.47 for the four parameter fit. Therefore, we selected the simpler, two parameter model to fit the performance data. **b** Same as in a for the BIC scores, with mean of 639.93 for the two parameter fit, 642.20 for the three parameter fit, and 643.73 for the four parameter fit. The lack of difference in the quality of the fits with or without the lapse rate parameters is consistent with the parameter estimation results of lapse rates in the hierarchical DDM (Extended Data Fig. 5i,j).

**Extended Data Fig. 3.**
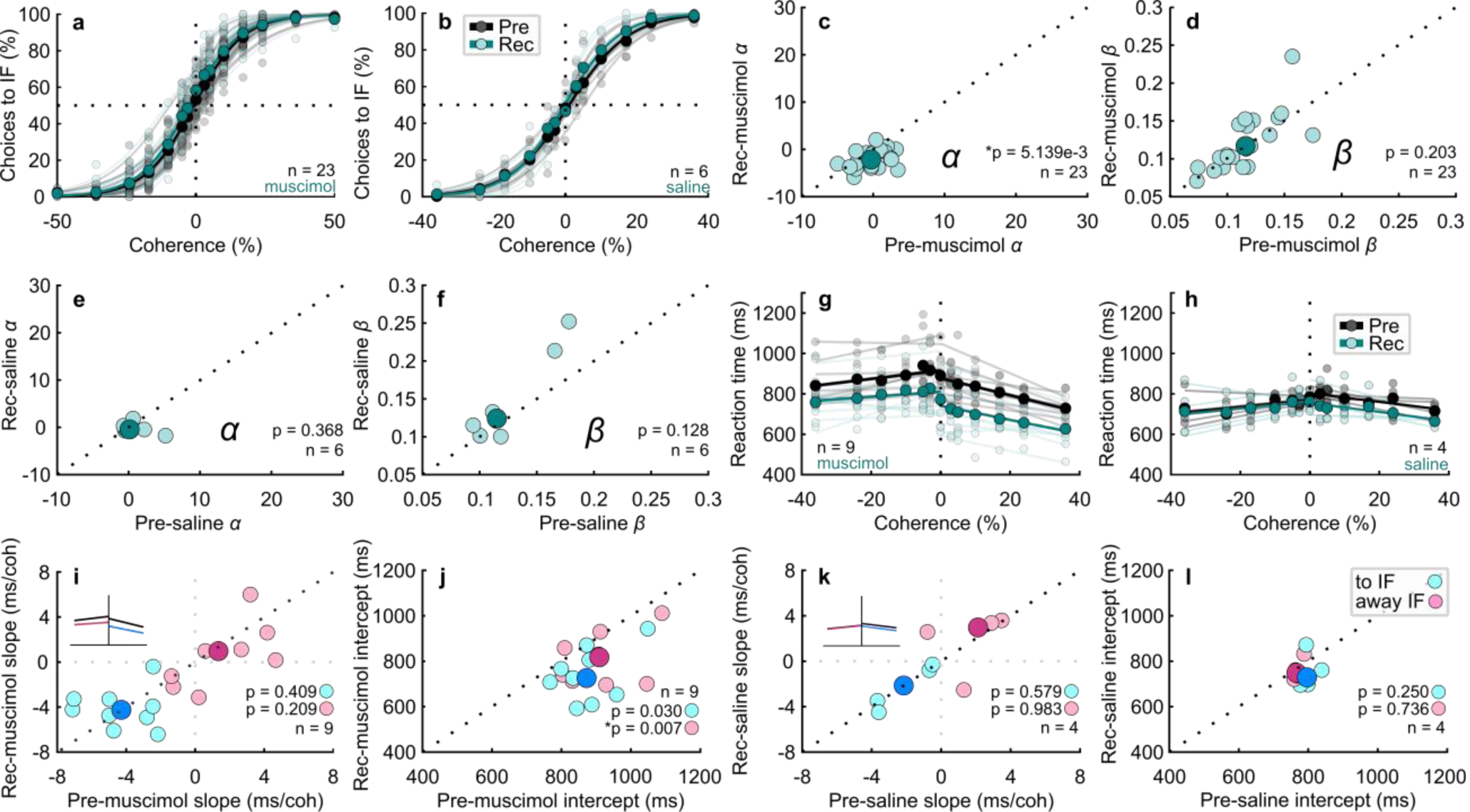
Decision-making behavior 24 hours after muscimol is similar to pre-muscimol decision-making behavior. (Associated with Fig. 2 of the main text) **a** Proportion of choices to the inactivated field (toIF) is plotted as a function of Glass pattern coherence. Black circles show pre-muscimol performance data and green circles show 24-hour recovery performance data. The black and green lines show the two parameter logistic fits to the performance data. n =23. **b** Same as in a for the pre-saline (black circles and lines) and the 24-hour recovery from saline (green circles and lines). n = 6. **c** *α* parameters from the logistic fits for the recovery data (Rec-muscimol) plotted against *α* parameters from the fits for the pre-muscimol data. On average, the *α* parameter shifted leftward during the recovery period compared to the pre-muscimol control (*w*(22) = 230, p = 0.005). Note that this was opposite to the direction of the shift that occurred post-muscimol as seen in the main Fig. 2a, as if the monkeys over-compensated for the effect of muscimol during recovery. **d** *β* parameters from the logistic fits for the recovery data plotted against the *β* parameters from the fits from the pre-muscimol data. On average, there were no significant differences in the *β* parameter (*t*(22) = -1.31, p = 0.20). **e-f** Same as in c and d for the saline experiments. **g** Reaction time (RT) plotted against coherence for the pre-muscimol data (black circles) and recovery data (green circles) from the RT version of the decision task (n = 9). The lines show linear fits to the RT data. The RT was shorter for the recovery data compared to the pre-muscimol data for all coherences. Similar to the results of the *α* parameter comparisons, the RT finding suggests a compensatory response to the muscimol injections 24 hours earlier. **h** Same as in g for the saline experiments. **i** The slope parameter from the linear fits to the RT data for the recovery data plotted against the pre-muscimol data. Cyan circles show the parameter of the linear fits of the RT data for toIF decisions (positive coherences) and magenta circles show the RT data for awayIF decisions (negative coherences). There were no significant differences on average (RT slope awayIF, *t*(8) = 1.37, p = 0.21; RT slope toIF, *t*(8) = -0.87, p = 0.41). **j** same as in i for the intercept parameter. There were significant changes in the intercept on average for the toIF side (RT intercept, *t*(8) = 3.61, p = 0.007) but not the awayIF side (RT intercept, *t*(8) =2.63, p = 0.03 n.s. Bonferroni correction). **k-l** Same as in i and j for the saline experiments. There were no significant differences in slope or intercept for these experiments (RT slope awayIF, *t*(3) = -0.02, p = 0.98; RT slope toIF, *t*(3) = 0.62, p = 0.58; RT intercept awayIF, *t*(3) = 0.37, p = 0.74; RT intercept toIF, *w*(3) = 9, p = 0.25). Note that four saline experiments were performed in the RT task and the other two were performed using the delayed version of the task so only four observations appear in this plot.

**Extended Data Table 2.** Statistics for recovery psychometric and chronometric function fits. (Associated with Fig. 2 of the main text) All statistical tests that were performed with the recovery data for the muscimol and saline injections for the *α, β*, RT slope, and RT intercept parameters associated with the fits in Extended Data Fig. 3 are listed. The first column indicates on which parameter the statistics were performed, the second column indicates which paired groups of data sessions were compared, the third column indicates which paired test was performed, the n indicates number of observations, followed by the p-value, test-statistic, and the confidence interval if a t-test was performed. The statistical results are sectioned by the injection (saline or muscimol) data that the statistics were performed on, marked by horizontal lines. We performed pairwise comparisons between parameters of pre-muscimol to recovery (the 2^nd^ pairwise comparison of pre-muscimol to post-muscimol are shown in the main text) with Bonferroni corrections (two pairwise comparison tests performed, α = 0.05/2 = 0.025).

| Muscimol psychometric function parameter comparisons |  |  |  |  |  |  |
| --- | --- | --- | --- | --- | --- | --- |
| Variables | Data Sessions | Test | n | p | Test stat. | CI |
| Alpha ($\alpha$) | Pre - rec | Wilcoxon signed rank test with Bonferroni correction | 23 | $5.139 \times 10^{-3}$ | 230 | — |
| Beta ($\beta$) | Pre - rec | Paired t-test with Bonferroni correction | 23 | 0.203 | -1.313 | (-0.019, 0.006) |
| Saline psychometric function parameter comparisons |  |  |  |  |  |  |
| Variables | Data Sessions | Test | n | p | Test stat. | CI |
| Alpha ($\alpha$) | Pre - rec | Paired t-test with Bonferroni correction | 6 | 0.368 | 0.988 | (-2.754, 5.257) |
| Beta ($\beta$) | Pre - rec | Paired t-test with Bonferroni correction | 6 | 0.128 | -1.822 | (-0.067, 0.018) |
| Muscimol chronometric function parameter comparisons |  |  |  |  |  |  |
| Variables | Data Sessions | Test | n | p | Test stat. | CI |
| Slope awayIF | Pre - rec | Paired t-test with Bonferroni correction | 9 | 0.209 | 1.366 | (-0.873, 2.593) |
| Slope toIF | Pre - rec | Paired t-test with Bonferroni correction | 9 | 0.409 | -0.872 | (-3.101, 1.609) |
| Intercept awayIF | Pre - rec | Paired t-test with Bonferroni correction | 9 | 0.030 | 2.630 | (-4.886, 216.526) |
| Intercept toIF | Pre - rec | Paired t-test with Bonferroni correction | 9 | 0.007 | 3.613 | (32.553, 240.499) |
| Saline chronometric function parameter comparisons |  |  |  |  |  |  |
| Variables | Data Sessions | Test | n | p | Test stat. | CI |
| Slope awayIF | Pre - rec | Paired t-test with Bonferroni correction | 4 | 0.983 | -0.023 | (-6.257, 6.189) |
| Slope toIF | Pre - rec | Paired t-test with Bonferroni correction | 4 | 0.579 | 0.621 | (-0.879, 1.186) |
| Intercept awayIF | Pre - rec | Paired t-test with Bonferroni correction | 4 | 0.736 | 0.370 | (-71.196, 85.050) |
| Intercept toIF | Pre - rec | Wilcoxon signed rank test with Bonferroni correction | 4 | 0.250 | 9 | — |

**Extended Data Table 3.**
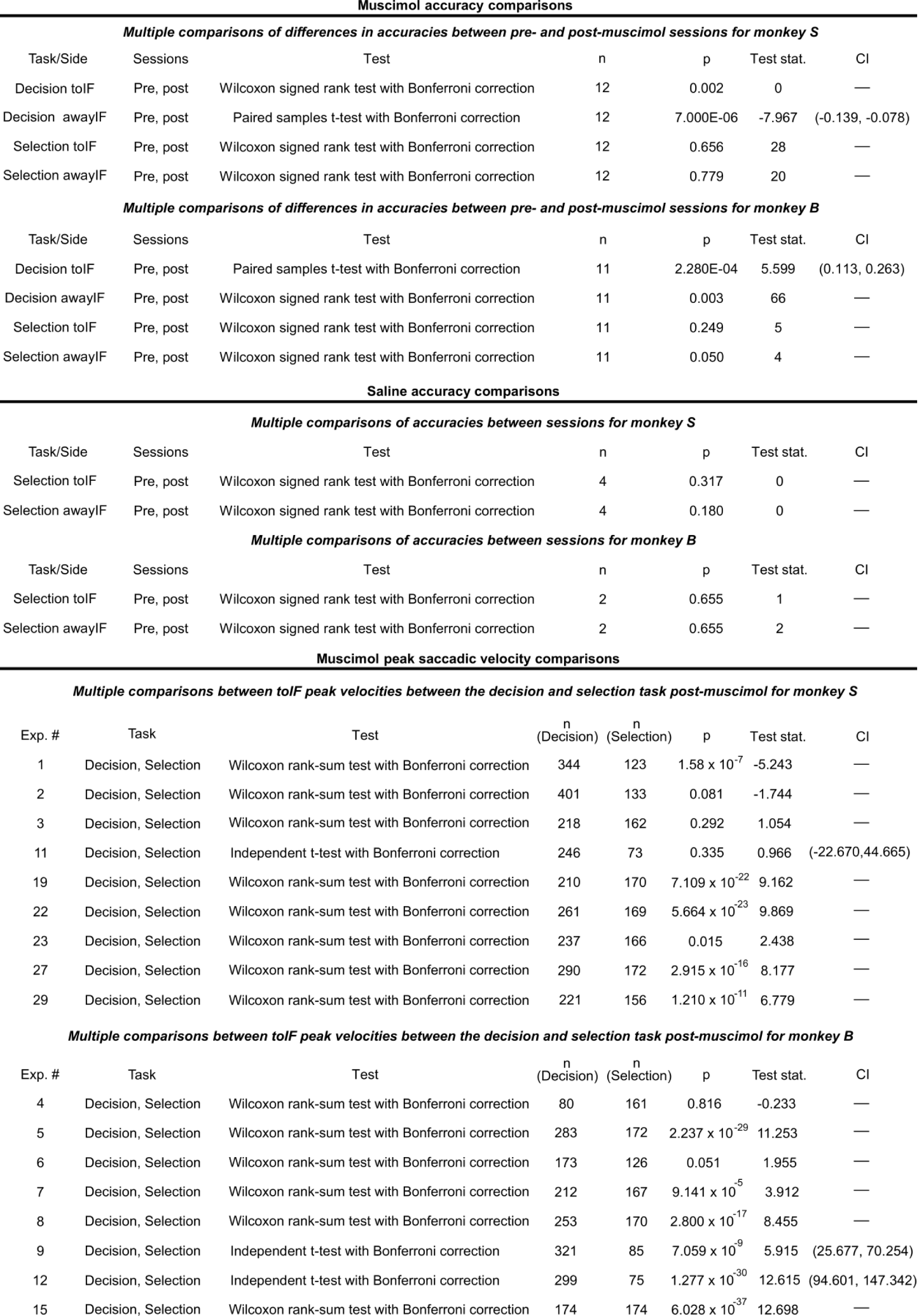
Statistics for choice accuracy and peak saccadic velocity analysis. (Associated with Fig. 3 of the main text) This table reports the statistical results of comparing differences in performance accuracy for the decision and selection tasks, for toIF and awayIF trials, pre- and post-muscimol. Also listed are the statistics for assessing the differences in post-muscimol peak saccadic velocity for toIF saccades made in the decision and the selection tasks, for each muscimol injection shown in Fig. 3d and e. N = 9 for monkey S and n = 8 for monkey B. The table is divided by horizontal black lines into three main sections - muscimol accuracy comparisons, saline accuracy comparisons, and muscimol peak saccadic velocity comparisons. Each of these main sections is further separated by monkey. The muscimol accuracy comparisons section and the saline accuracy comparisons section have the same ordering of columns. The first column, “task/side” describes the task and side of accuracy from pre to post, i.e. “decision toIF” means the test assessed differences in accuracy for toIF trials in the decision task between pre and post-muscimol. The second column describes the data sessions compared and the next column describes the statistical test that was used to compare the two data sets. We used a paired samples t-test when both samples (data from pre and post) consisted of normally distributed data or the Wilcoxon signed rank test when either of the two samples consisted of non-normally distributed data, implementing a Bonferroni correction (four accuracy multiple comparison tests per monkey, *α* = 0.05/4 = 0.0127). The next column indicates the sample size (number of injections). The column after shows the p-value of the statistical comparison analysis followed by the test statistic. The last column indicates the confidence interval if a t-test was performed. For the section assessing differences in post-muscimol peak saccadic velocity for toIF saccades made in the decision task compared to the selection task, on an injection-by-injection basis, the first column describes the injection experiment number and the second column describes the specific statistical test that was used to compare the two samples of peak velocities of saccades made. A two independent sample t-test (for normally distributed data sessions) or Wilcoxon rank-sum test (for non-normally distributed data sessions) was used to compare the post-muscimol toIF peak saccadic velocities between the decision and selection tasks with Bonferroni corrections of *α* = 0.05/9 = 0.0056 for the tests for monkey S, since nine t-tests were performed on data from each injection, and *α* = 0.05/8 = 0.0063 for monkey B, since eight t-tests were performed on data from each injection, totaling 17 injections. Six injections were excluded due to technical issues with the eye tracker that impacted measurement of eye speed but not assessment of choice or RT. The next column indicates the sample size, the number of trials where the peak saccadic velocities were recorded, from the decision task and the following column indicates the sample size of the peak saccadic velocities from the selection task. The next column relates the p-value of the test followed by the test statistic in the next column for the specific statistical test performed. The last column indicates the confidence interval if a t-test was performed.

| Muscimol accuracy comparisons |  |  |  |  |  |  |  |
| --- | --- | --- | --- | --- | --- | --- | --- |
| Multiple comparisons of differences in accuracies between pre- and post-muscimol sessions for monkey S |  |  |  |  |  |  |  |
| Task/Side | Sessions | Test | n | p | Test stat. | CI |  |
| Decision toIF | Pre, post | Wilcoxon signed rank test with Bonferroni correction | 12 | 0.002 | 0 | — |  |
| Decision awayIF | Pre, post | Paired samples t-test with Bonferroni correction | 12 | 7.000E-06 | -7.967 | (-0.139, -0.078) |  |
| Selection toIF | Pre, post | Wilcoxon signed rank test with Bonferroni correction | 12 | 0.656 | 28 | — |  |
| Selection awayIF | Pre, post | Wilcoxon signed rank test with Bonferroni correction | 12 | 0.779 | 20 | — |  |
| Multiple comparisons of differences in accuracies between pre- and post-muscimol sessions for monkey B |  |  |  |  |  |  |  |
| Task/Side | Sessions | Test | n | p | Test stat. | CI |  |
| Decision toIF | Pre, post | Paired samples t-test with Bonferroni correction | 11 | 2.280E-04 | 5.599 | (0.113, 0.263) |  |
| Decision awayIF | Pre, post | Wilcoxon signed rank test with Bonferroni correction | 11 | 0.003 | 66 | — |  |
| Selection toIF | Pre, post | Wilcoxon signed rank test with Bonferroni correction | 11 | 0.249 | 5 | — |  |
| Selection awayIF | Pre, post | Wilcoxon signed rank test with Bonferroni correction | 11 | 0.050 | 4 | — |  |
| Saline accuracy comparisons |  |  |  |  |  |  |  |
| Multiple comparisons of accuracies between sessions for monkey S |  |  |  |  |  |  |  |
| Task/Side | Sessions | Test | n | p | Test stat. | CI |  |
| Selection toIF | Pre, post | Wilcoxon signed rank test with Bonferroni correction | 4 | 0.317 | 0 | — |  |
| Selection awayIF | Pre, post | Wilcoxon signed rank test with Bonferroni correction | 4 | 0.180 | 0 | — |  |
| Multiple comparisons of accuracies between sessions for monkey B |  |  |  |  |  |  |  |
| Task/Side | Sessions | Test | n | p | Test stat. | CI |  |
| Selection toIF | Pre, post | Wilcoxon signed rank test with Bonferroni correction | 2 | 0.655 | 1 | — |  |
| Selection awayIF | Pre, post | Wilcoxon signed rank test with Bonferroni correction | 2 | 0.655 | 2 | — |  |
| Muscimol peak saccadic velocity comparisons |  |  |  |  |  |  |  |
| Multiple comparisons between toIF peak velocities between the decision and selection task post-muscimol for monkey S |  |  |  |  |  |  |  |
| Exp. # | Task | Test | n<br>(Decision) | n<br>(Selection) | p | Test stat. | CI |
| 1 | Decision, Selection | Wilcoxon rank-sum test with Bonferroni correction | 344 | 123 | 1.58 x 10<sup>-7</sup> | -5.243 | — |
| 2 | Decision, Selection | Wilcoxon rank-sum test with Bonferroni correction | 401 | 133 | 0.081 | -1.744 | — |
| 3 | Decision, Selection | Wilcoxon rank-sum test with Bonferroni correction | 218 | 162 | 0.292 | 1.054 | — |
| 11 | Decision, Selection | Independent t-test with Bonferroni correction | 246 | 73 | 0.335 | 0.966 | (-22.670,44.665) |
| 19 | Decision, Selection | Wilcoxon rank-sum test with Bonferroni correction | 210 | 170 | 7.109 x 10<sup>-22</sup> | 9.162 | — |
| 22 | Decision, Selection | Wilcoxon rank-sum test with Bonferroni correction | 261 | 169 | 5.664 x 10<sup>-23</sup> | 9.869 | — |
| 23 | Decision, Selection | Wilcoxon rank-sum test with Bonferroni correction | 237 | 166 | 0.015 | 2.438 | — |
| 27 | Decision, Selection | Wilcoxon rank-sum test with Bonferroni correction | 290 | 172 | 2.915 x 10<sup>-16</sup> | 8.177 | — |
| 29 | Decision, Selection | Wilcoxon rank-sum test with Bonferroni correction | 221 | 156 | 1.210 x 10<sup>-11</sup> | 6.779 | — |
| Multiple comparisons between toIF peak velocities between the decision and selection task post-muscimol for monkey B |  |  |  |  |  |  |  |
| Exp. # | Task | Test | n<br>(Decision) | n<br>(Selection) | p | Test stat. | CI |
| 4 | Decision, Selection | Wilcoxon rank-sum test with Bonferroni correction | 80 | 161 | 0.816 | -0.233 | — |
| 5 | Decision, Selection | Wilcoxon rank-sum test with Bonferroni correction | 283 | 172 | 2.237 x 10<sup>-29</sup> | 11.253 | — |
| 6 | Decision, Selection | Wilcoxon rank-sum test with Bonferroni correction | 173 | 126 | 0.051 | 1.955 | — |
| 7 | Decision, Selection | Wilcoxon rank-sum test with Bonferroni correction | 212 | 167 | 9.141 x 10<sup>-5</sup> | 3.912 | — |
| 8 | Decision, Selection | Wilcoxon rank-sum test with Bonferroni correction | 253 | 170 | 2.800 x 10<sup>-17</sup> | 8.455 | — |
| 9 | Decision, Selection | Independent t-test with Bonferroni correction | 321 | 85 | 7.059 x 10<sup>-9</sup> | 5.915 | (25.677, 70.254) |
| 12 | Decision, Selection | Independent t-test with Bonferroni correction | 299 | 75 | 1.277 x 10<sup>-30</sup> | 12.615 | (94.601, 147.342) |
| 15 | Decision, Selection | Wilcoxon rank-sum test with Bonferroni correction | 174 | 174 | 6.028 x 10<sup>-37</sup> | 12.698 | — |

**Extended Data Fig. 4.**
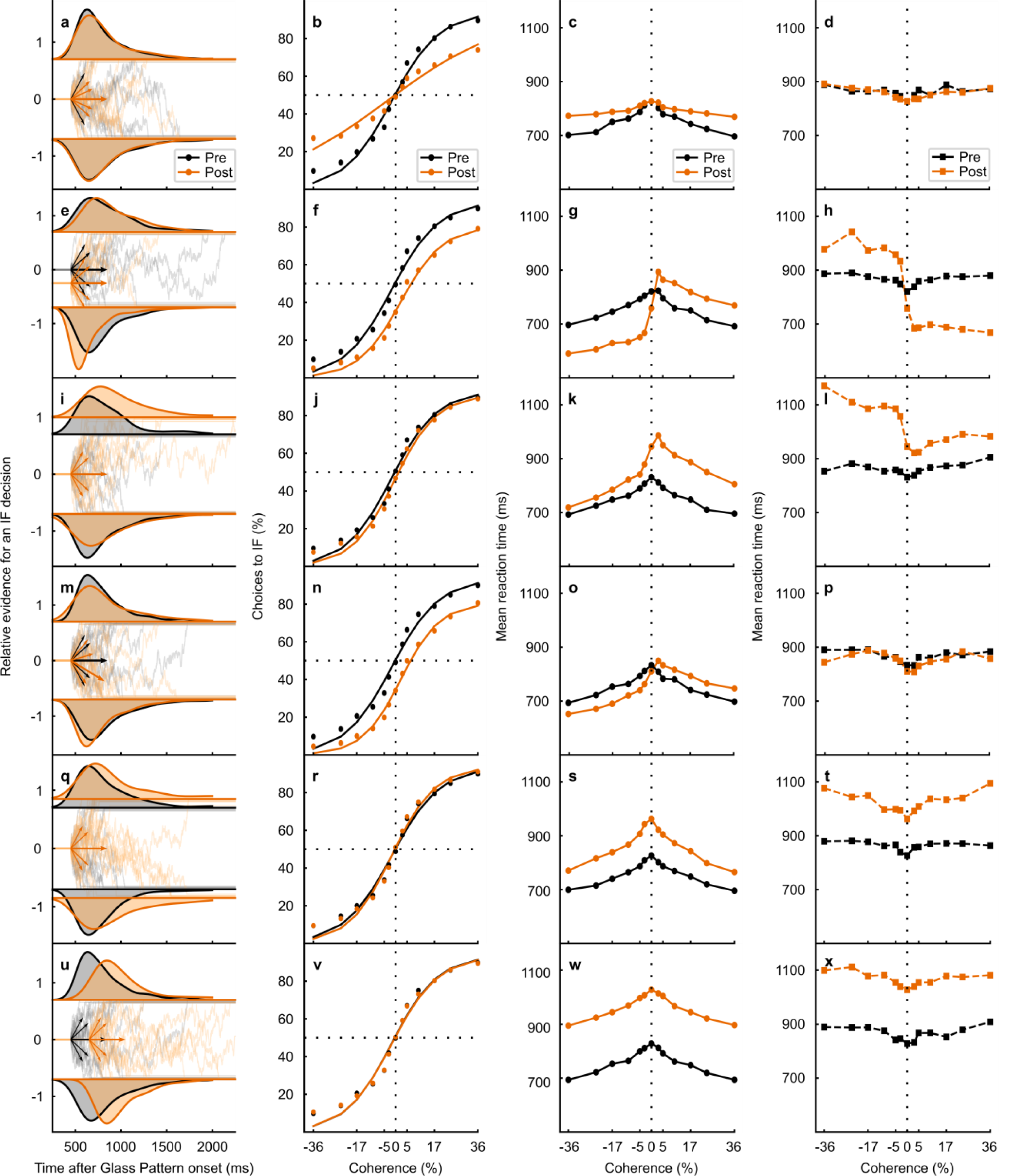
DDM model simulations for changes in model parameters. (Associated with Fig. 4 of the main text). Panels a-p are the same as those shown in Fig. 4 of the main text. **a** RT distribution from the 0% coherence condition (density approximated through kernel smoothing) predicted by a DDM simulation with only a multiplicative decrease (gain change) in drift rates for post-muscimol (orange). Pre-muscimol shown in black. Below the RT distributions, the relative evidence for toIF decisions is plotted over time since the Glass pattern onset and the short arrows show drift rates for toIF decisions (positive) and awayIF decisions (negative) pre- and post-muscimol, for the 0%, 10%, and 36% coherence conditions. The longer arrows show the mean drift rate across coherences (the mean drift rates over all toIF and awayIF coherences^15^). **b** The psychometric function, plotted as a proportion of toIF choices over coherences, predicted by the DDM variant simulation with a change in the multiplicative decrease (gain decrease) in drift rates, which changes the slope (without a shift) of the psychometric function. A change in the slope of the psychometric function was not observed in the data (Fig. 4r, v), making the multiplicative change in drift rates (a gain change), an unlikely explanation for the observed data. **c** Mean RT predictions for correct trials for each coherence condition for the DDM simulation with a multiplicative decrease in drift rates for pre (black) and post (orange). **d** Same as in c but for error trials. **e-h** Same as in a-d but for the DDM variant simulation with only a change in proportional start-point of the evidence accumulation path away from the IF (often interpreted as an initial bias away from the IF). A decrease in proportional start point away from the IF predicts a shift in the psychometric function as observed in the real data (Fig. 4r,v), making a change in the proportional start point a possibility in explaining the decision bias we observed in the post-muscimol data. However, a start point change away from the IF also predicts a decrease in error toIF RTs which we did not observe in the data (Fig. 4t,x). **i-l** Same as in a-d but for the DDM variant with an increase in the upper boundary but no absolute start point change (start point proportionally decreased away from the IF). This parameter change also predicts a shift in the psychometric function away from IF decisions as we observed in the data (Fig. 4r,v). However, this parameter change cannot explain the magnitude of the psychometric function shift we observed (Fig. 4r,v) with similar changes in simulated and observed mean RTs (Fig. 4s,t,w,x). **m-p** Same as in a-d but for the DDM variant with only the same additive shift in all drift rates away from the IF, representing a change in the mean drift rate across coherences. The psychometric function predictions of the model simulation with a change in the mean drift rate across coherences predict a shift in the psychometric function that is observed in the data (Fig. 4r, v). The increases in correct mean RT for awayIF decisions are predicted and shown for both monkeys (Fig. 4s, w). Overall, a change in mean drift rate across all coherences (drat rate criterion) is mostly likely to explain the data we obtained after muscimol inactivation of the SC. **q-t** Same as in a-c but for the DDM model variant that describes RT distributions and performance with only a large increase in the symmetric boundaries. This parameter change predicts only slight steepening of the slope of the psychometric function and no changes in the shift of the psychometric function as observed in the data (Fig. 4r,v), making the symmetric boundary change an unlikely possibility for explaining the effects of SC inactivation. **u-x** Same as in a-c but for the DDM variant that describes RT distributions and performance with only an increase in non-decision time. Non-decision time changes do not explain any changes in performance and thus cannot explain a shift in the psychometric function observed in the data from both monkeys (Fig. 4r,v), making a change in non-decision time unlikely to explain the effects of SC inactivation on decision-making.

**Extended Data Fig. 5.**
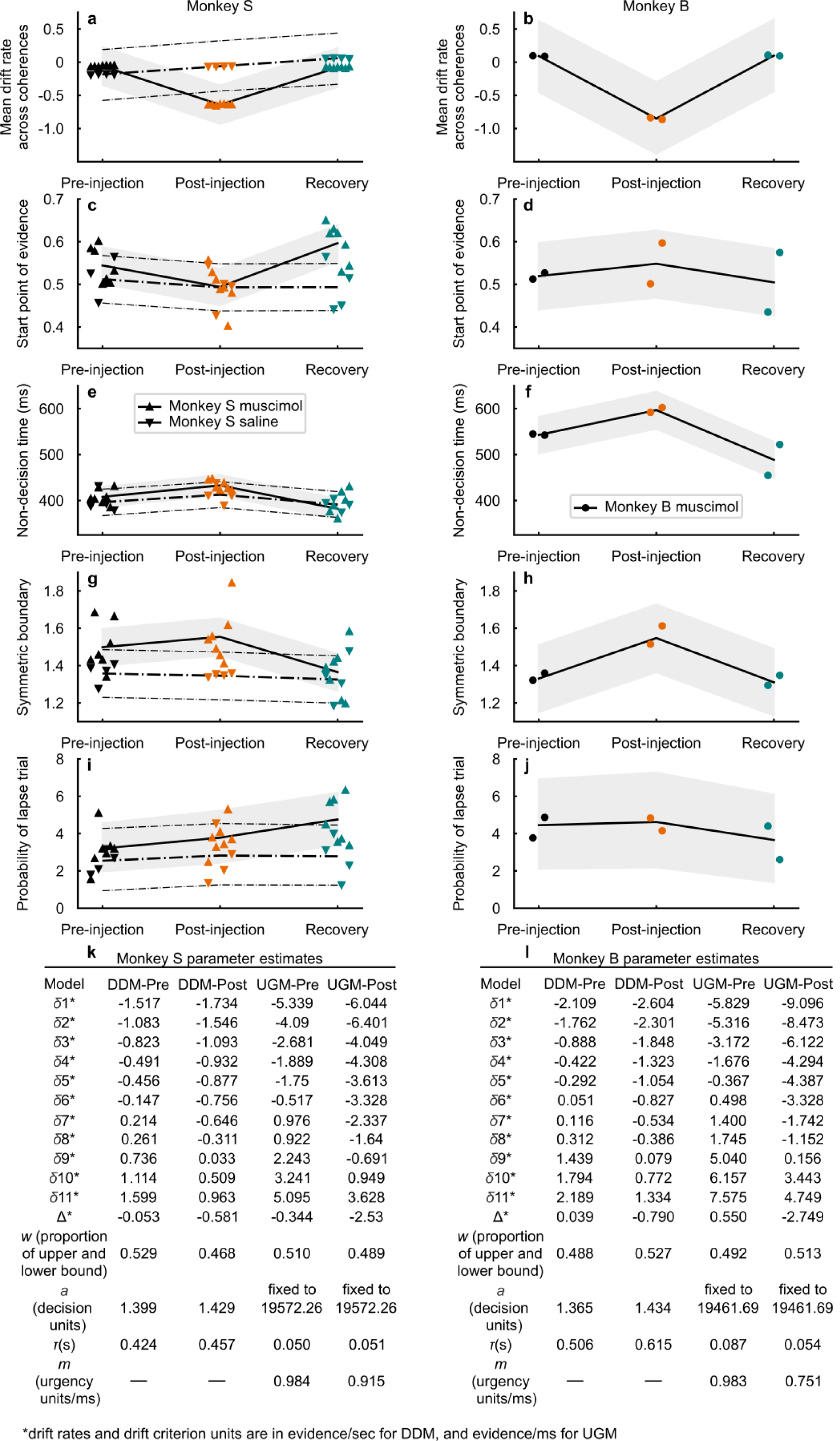
Parameter estimates for HDDM, DDM, and UGM (Associated with Fig. 4 of the main text). **a-j** Estimates from the full HDDM of hierarchical parameters (*μ*) for each monkey (solid lines in the muscimol experimental condition; dotted lines for monkey S in the saline experimental condition, we did not collect data from the RT task for monkey B in the saline condition). 95% credible intervals with 2.5th and 97.5th quantile boundaries of hierarchical parameters provided by shading for the muscimol condition and smaller dot-dashed lines for the saline condition. Also shown are individual session parameter estimates for monkey S’s muscimol data (upward-pointing triangles), monkey B’s muscimol data (circles), and monkey S’s saline data (downward-pointing triangles). Estimates were obtained from the median posterior distributions of each parameter. **a** Estimates of the HDDM session-level mean drift rate (*Δ*) and hierarchical mean drift rate (*μ*_*Δ*_) for monkey S (pre BF = 0.08, post BF = 3.19 × 10^6^, 99.7% probability of decrease pre to post). **b** Same as in a but for monkey B (pre Bayes factor BF = 0.14, post BF = 17.87, 99.0% probability of decrease pre to post). **c** Estimates of the HDDM session-level start point (*w*) and hierarchical start point (*μ*_*w*_) for monkey S (pre BF=0.73, post BF = 0.09, 95.1% probability of a proportional start point bias away from the IF from pre to post). **d** Same as in c but for monkey B (pre BF = 0.18, post BF = 0.35, 70.6% probability of a proportional start point bias towards the IF from pre to post). **e** Estimates of the session-level non-decision time (*τ*) and hierarchical non-decision time (*μ*_*τ*_) for monkey S (94.5% probability of an increase from pre to post). **f** Same as in e for monkey B (97.0% probability of increase pre to post). **g** Estimates of the session-level symmetric boundary (*a*) and hierarchical symmetric boundary (*μ*_*a*_) for monkey S (78.9% probability of an increase from pre to post). **h** Same as in g but for monkey B (95.2% probability of increase pre to post). **i** Estimates of the session-level lapse proportion (*λ*) and hierarchical lapse proportion (*μ*_*λ*_) for monkey S (72.5% probability of increase from pre to post). **j** Same as in i but for monkey B (54.0% probability of increase pre to post). **k-l** The parameter estimates obtained from fitting the DDM and the UGM to the pre- and post-muscimol data for monkey S (panel k) and monkey B (panel l). The first row describes the model that was fit (DDM or UGM) and which data session (pre or post) was used to fit the model. The next 11 rows represent the drift rate parameter estimates (*δ*_*k*_), in evidence units/sec for the DDM or evidence units/ms for the UGM, for the *k* = 11 conditions (−24%, -17%, -10%, -3%, -5%, 0%, 5%, 3%, 10%, 17%, 24% coherences). The next row shows the mean drift rate collapsed across coherence (*Δ*). This parameter was not explicitly fit, but rather calculated from the drift rates that were estimated from fits to summarize the change in drift rates. The mean drift rate across coherences decreased from pre- to post-muscimol for both DDM and UGM and for both monkeys (difference in monkey S, 0.53 evidence units/sec decrease for DDM, 2.19 evidence units/ms decrease for UGM; monkey B, 0.83 evidence units/sec decrease for DDM, 3.30 evidence units/ms decrease for UGM). The next row shows the proportional start point parameter *w*, defined as the proportion of the distance between the upper and lower bound. For monkey S, the start point parameter had slightly decreased from pre- to post-muscimol in both the DDM (0.06 decrease) and UGM (0.02 decrease), indicating the start point moved closer to the awayIF decision bound, and for monkey B, the start point parameter slightly increased in the DDM (0.04 increase) and UGM (0.02 increase), indicating the start point moved closer to the toIF decision bound. The next row shows the bound height parameter *a*, defined as the distance between the upper and lower bounds. For both monkeys, but more prominent in monkey B, the bound parameter had slightly increased from pre to post in the DDM (monkey S, pre to post increase of 0.03 decision units; monkey B, pre- to post-muscimol increase of 0.07 decision units), whereas the bound was fixed in the UGM (Supplemental Methods). The row after shows the non-decision time *τ*, in seconds, where we see a slight increase in the DDM (0.03 sec increase) and UGM (0.001 sec increase) for monkey S and a greater increase in the DDM for monkey B (0.11 sec increase), but not for the UGM (0.03 sec decrease). The last row shows the urgency slope estimates for the UGM, *m*, decreasing slightly with muscimol for monkey S (0.07 urgency units/ms), and decreasing more for monkey B (0.23 urgency units/ms).

**Extended Data Table 4.**
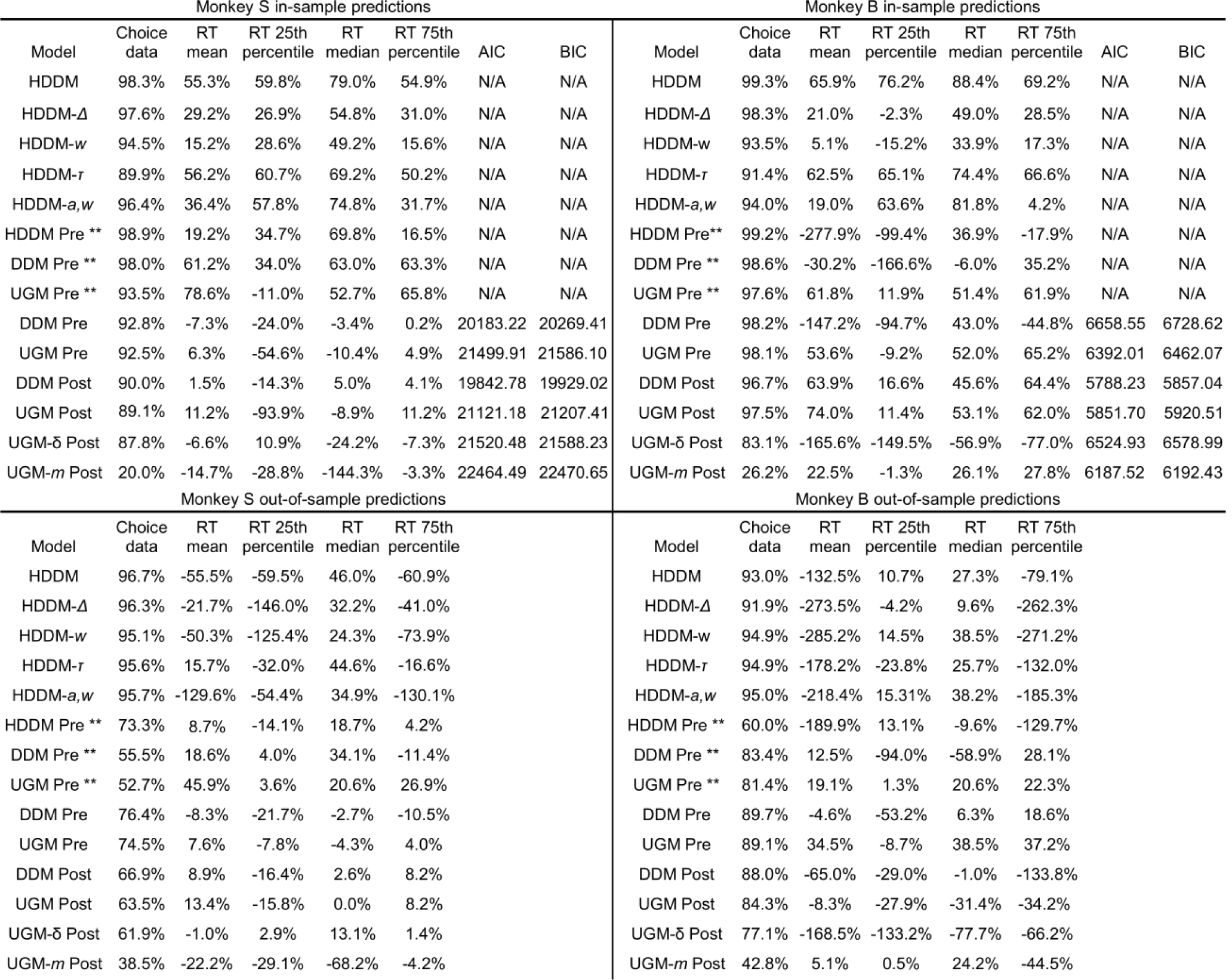
Table of *R*^*2*^_*pred*_ for in sample and out of sample predictions. (Associated with Fig. 4 of the main text) For model comparisons, we obtained goodness of fit measures such as *R*^*2*^_*pred*_ for choice performance data, RT mean, RT 25th percentile, RT 75th percentiles, along with the AIC and BIC values (where can be applied) for the in-sample predictions where various models were fit to a random 80% of the data (Supplemental Methods) for monkey S (top left) and monkey B (top right). Also shown are the *R*^*2*^_*pred*_ values for out-of-sample predictions (bottom left for monkey S and bottom right for monkey B), where the predictions of the fits are compared to the 20% of the data that was not fit to assess if the data was overfit and see if the same conclusions of model comparison are supported. Each section of the table starts with the model comparisons between the full HDDM, HDDM with free mean drift rate across coherences (HDDM-*Δ*), HDDM with free proportional start point (HDDM-*w*), the HDDM with free non-decision time (HDDM-*τ*), and the HDDM with the free proportional start point and bound (HDDM-*a,w*). For both monkeys, the in-sample *R*^*2*^_*pred*_ shows that the HDDM-*Δ* fit explains the choice data best compared to HDDM variants with specific parameters free to vary. The HDDM*-a,w* also captures some of the choice data but not to the extent of the HDDM-*Δ*, which fully captures the magnitude of the decision bias, which is visualized in Extended Data Fig. 6. Like HDDM*-a,w*, the HDDM-*w* also captures some of the decision bias, but not sufficiently to capture the magnitude of decision bias seen in the data. The HDDM*-τ* explains the least of the choice data out of the models. The predictions from the model fits denoted with ** are the models that were fit to individual pre-muscimol data (rather than pooled pre-muscimol data) with none conditions (17%, 10%, 5%, 3% separately for toIF and awayIF trials, with the 0% collapsed across toIF and awayIF trials) for direct model comparison between the HDDM, DDM, and UGM for each monkey (Supplemental methods) to see whether the UGM is also a reasonable decision-making model assumption as the HDDM/DDM in our behavioral data. The next four models are the DDM and UGM fits for pre and post data that were pooled (Supplemental Methods) to show the measures of fits that correspond to the parameter estimates seen in Extended Data Fig. 5. Finally, the model comparisons for the full UGM, UGM with free drift rates (UGM-*δ*), and UGM with free slope (UGM-*m*) for the post-muscimol data (while keeping the other parameters fixed at pre-muscimol parameter estimate values, Supplemental Methods), are shown. For monkey S, the AIC and BIC values indicate that the UGM-*δ* is a better fit than the UGM-*m*, whereas for monkey B, the AIC and BIC values favor the UGM-*m* than the UGM-*δ*. However, the in-sample *R*^*2*^_*pred*_ values for choice data show that the UGM-*m* fails to capture the choice data for both monkeys whereas the UGM-*δ* captures the choice data almost as well as the full UGM (also visualized in Extended Data Fig. 6). The UGM-*δ* better explains the shift in decision bias than UGM-*m* in both monkeys, while the UGM-*m* better explains the RT distribution change from monkey B, but not for monkey S, likely due to individual strategy differences. The out-of-sample predictions generally show the same results although they are more variable.

**Extended Data Fig. 6.**
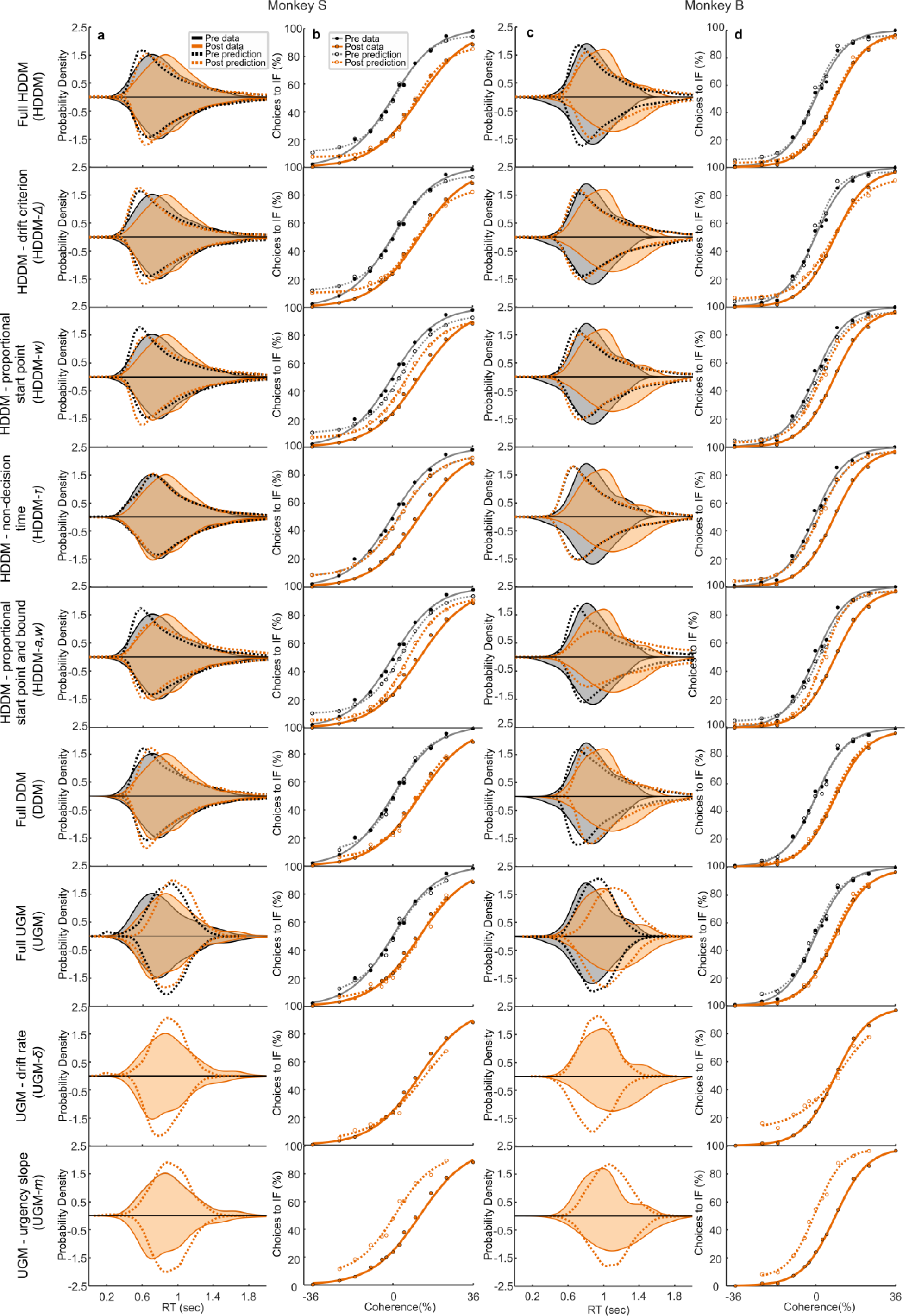
Model predictions versus data for RT distributions and psychometric functions. (Associated with Fig.4 of the main text). Column **a** shows the predicted RT distributions (0% coherence, density approximated through kernel smoothing) from the DDM, HDDM and UGM model variants (dashed lines) together with the actual data (sold lines), pre-muscimol (black) and post-muscimol (orange), for monkey S. We observed a rightward skew of the RT distribution, consistent with a fixed bound model of decision-making and captured by the DDM rather than the UGM as was also indicated by the *R*^*2*^_*pred*_, AIC, and BIC goodness of fit values (Extended Data Table 4). Column **b** shows the same as in a but for psychometric functions (performance data, four parameter logistic model using equation shown in Extended Data Fig. 2). Column **c** shows the same as in a for monkey B’s data and model fits. The RT distributions from monkey B were more normally distributed compared to the skewed RT distributions of monkey S, suggesting that the UGM rather than the DDM would explain monkey B’s data, consistent with the goodness of fit values (Extended Data Table 4). Column **d** shows the same as in c but for psychometric function (performance data). Each row indicates the results of each model’s prediction compared to data for both monkey S and monkey B. The models from top to bottom are the full HDDM, HDDM with a free-to-vary mean drift rate across coherences (HDDM-*Δ*), HDDM with a free-to-vary proportional start point (HDDM-*w*), HDDM with free-to-vary non-decision time (HDDM-*τ*), HDDM with both a proportional start point and bound free to vary (HDDM-*a,w*), the non-hierarchical DDM, the full UGM, the UGM with free-to-vary drift rates (UGM-*δ*), and UGM with a free-to-vary urgency slope (UGM-*m*). Note that only the post-muscimol data are shown for the UGM with a single free parameter since we only fit the post data with those models where we let only one parameter free to vary while the rest of the parameters were fixed to pre-muscimol parameter estimates (Supplemental Methods). Also for the DDM and UGM fits, note that there are only 11 conditions (−24 to 24 % coherence) for the psychometric functions because only 11 conditions were fit (Supplemental Methods). For the HDDM, out of all the variants (first five rows), the full HDDM predictions visually match the data for both performance and RT. The the prediction of the HDDM-*Δ* captures the decision bias from the data almost equally well for both monkeys. The prediction for the HDDM-*w* and HDDM-*a,w* also predicts a decision bias, but is insufficient to explain the magnitude of the shift in decision bias that we observed in the data. The HDDM-*τ* fails to capture any decision bias (RT and performance predictions for pre and post are overlapping). The predictions of the simple DDM also capture the 0% RT distribution well, more so for monkey S than for monkey B, and also capture the choice data well. The opposite is true for the full UGM predictions, where the RT predictions capture monkey B’s data more than monkey S (see goodness of fit values in Extended Data Table 4), but also captures performance data well for both monkeys. The UGM-*δ* captures the shift in decision bias from the post-muscimol data from both monkeys, consistent with the findings from the HDDM, whereas the UGM-*m* fails to capture the decision bias in the post data.

## SUPPLEMENTARY METHODS

We undertook a multipronged analytical approach to assess the effects of unilateral superior colliculus (SC) inactivation on perceptual decision-making in monkeys. First, we evaluated changes in psychometric functions and mean RTs pre- and post-muscimol for decisions to and away from the inactivated field (IF; Fig. 2). Second, we evaluated changes in the shapes of RT distributions pre- and post-muscimol for decisions to and away from the IF as the shapes of RT distributions provide reliable ways to distinguish changes in different decision processes^31^ (Fig. 4 and Extended Data Fig. 6). Third, we simulated DDMs with changes only in specific parameters from pre- to post-muscimol to assess and compare predicted changes in psychophysical performance and mean RTs that may occur if different aspects of decision-making were changed post-muscimol (Fig.4, Extended Data Fig. 4). Fourth, we fit a hierarchical drift-diffusion model, HDDM, to the data and estimated parameters to see what parameters actually changed due to the effect of SC inactivation (Extended Data Fig. 5). Fifth, to determine which parameter best explained the changes observed with muscimol in the SC, we fit HDDMs to the data allowing different model parameters to vary individually and compared these to HDDM with all parameters allowed to vary to determine which model best explained the results of the muscimol inactivation (Extended data Table 4 and Extended data Fig. 6). We also fit a non-hierarchical DDM, therefore using two different methods of parameter estimation (hierarchical Bayesian estimation and Quantile Maximum Probability Estimation, QMPE) to ensure robustness (Extended Data Fig. 5). Finally, to determine whether an altered urgency signal explained the effects of unilateral inactivation of the SC during decision-making, we also fit UGMs to the pre- and post-muscimol data to see which parameters changed due to the muscimol, and then compared to UGMs with urgency or drift rate parameters free to vary to determine which model best explained the post-muscimol data (Extended data Table 4 and Extended data Fig. 6).

### Signal Detection Theory (SDT) analysis

In addition to the logistic fits as described in the Methods and shown in Fig. 2, we also calculated *d’* and *c* to assess whether unilateral inactivation of the SC with muscimol impaired perceptual sensitivity or decision criterion, respectively^17^. To calculate *d*’, we used the following equation modified from Macmillian^30^;

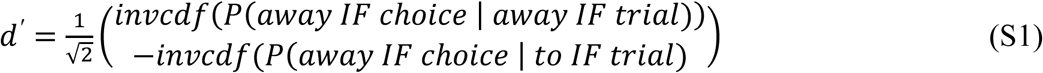

To calculate the position of the decision criterion we also used the equation from Macmillan^30^;

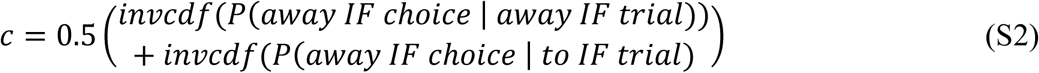

Extended data Fig. 2 shows the *d’* and *c* results for both monkeys. Overall, there was a significant change in criterion after muscimol that caused a greater decision bias away from the IF, whereas sensitivity, *d’*, showed no significant change after muscimol. Neither parameter changed after saline injections.

### Non-parametric reaction time (RT) analysis

To visualize and analyze the chronometric functions from the reaction time (RT) version of the decision task (seven in monkey S and two in monkey B for muscimol injections, and four in monkey S for saline injections), we used least squares regression to fit the RT data with a linear function of the form:

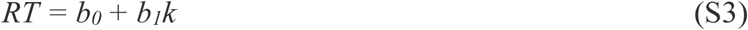

This was fit separately for the positive (toIF) and negative (awayIF) coherences (*k*). We extracted the slope parameter (*b*_*1*_), to determine coherence dependent changes in RT from pre- to post-muscimol and recovery, and the intercept parameter (*b*_*0*_), to assess general, non-coherence dependent changes in RT from pre- to post-muscimol and recovery. The pre- and post-muscimol data and results appear in Fig. 2 and the recovery data appear in Extended data Fig. 3. Coherence dependent changes occurred to toIF decisions and not for awayIF decisions.

### Drift-diffusion model (DDM) simulations

DDMs assume that decision-makers accumulate a representation of noisy sensory evidence over time. An evidence path on one trial reflects an internal representation of orientation evidence obtained by visually sampling the Glass Pattern until a fixed amount of evidence is reached for a decision, reported by a saccade, to or away from the IF. Different aspects of the decision-making processes are reflected in different parameters of the DDM as described in the Methods^31^. The DDM process was defined and simulated by the following equations:

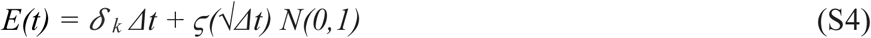

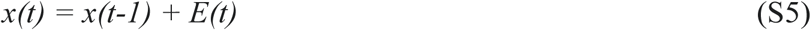

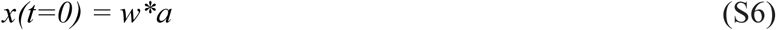

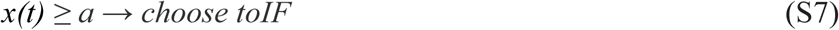

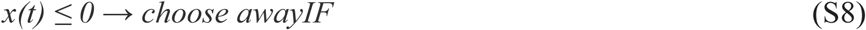

The evidence at time (*t*), denoted by *E(t)*, depends on the drift rate (*δ*_*k*_) for each condition (*k*) with an accumulation noise (*ϛ*), for every infinitely small time step *Δt*. In simulations of DDMs and in model fitting using QMPE (discussed later), the small time step is estimated with *Δt = 1* ms and the noise constant is set to *s=1* evidence unit. Evidence is accumulated during a trial until one of two boundaries are reached (*a* or 0), marking the decision variable *x(t*). The proportional start point parameter (*w*) is defined as the proportion of the distance between the two bounds (*a*) such that *w* = 0.5 indicates the start point is in the midpoint between the upper (toIF) and lower (awayIF) bound, *w* > 0.5 indicates the start point is closer to the upper (toIF) bound, and *w* < 0.5 indicates the start point is closer to the lower (awayIF) bound.

There are at least six possible, non-mutually exclusive ways that unilateral SC inactivation could alter decision-making, which we show in simulation using the DDM (Fig. 4, Extended Data Fig. 4). These simulations yielded different expectations for the psychometric and chronometric functions of the data depending upon how SC inactivation impacts decision-making. Five hundred trials were simulated for each possible scenario to make the predictions of choice performance and RT distributions, and the first 20 trial simulations of the evidence paths are shown in Fig. 4 and Extended Data Fig. 4. All these simulations of performance and RT data can be used for visual comparison with the actual data obtained from monkey S and B shown in Fig 4q-x.

The first possibility is that muscimol impairs or alters the drift rate (*δ*) in a way that changes it’s the slope over coherence values (i.e., a multiplicative decrease or *gain* effect). The absolute change in drift rates across coherence values are determined by the internal strength of the sensory evidence for orientation during the decision task. This change in absolute slope in drift rates could be considered analogous to a change of sensitivity in SDT (in addition to a symmetric change in boundary *a*, discussed below). We simulated a DDM (Fig. 4a-d, Extended Data Fig. 4a-d) with only a multiplicative decrease in drift rates of 0.5 (pre-muscimol *δ*_*1k*_ spanned -3 to 3 with steps of 0.5 away with mean *Δ*_*1*_ = 0 and post-muscimol *δ*_*2k*_ spanned -1.5 to 1.5 with steps of 0.25 away with mean *Δ*_*2*_ = 0) and no other parameter changes (*a* = 1.4, *τ* = .45 sec, *w* = 0.5, *λ* = 0). Fig. 4a and Extended Data Fig. 4a shows the representation of a sequential sampling of relative evidence towards one of two boundaries with 20 simulated evidence paths for a simulated pre-muscimol injection condition by thin grey lines and a simulated post-muscimol injection condition by faded orange lines in the 0% coherence condition. A choice is made once the relative path passes one of two boundaries, then a saccade to the IF or saccade away from the IF occurs. Simulated RT distributions from the 0% coherence condition are shown on their respective boundaries for each choice (Fig. 4a, Extended Data Fig. 4a; shaded orange RT curves from the post-muscimol condition in front of shaded grey RT curves from the pre-muscimol condition). Drift rates are the mean rate of evidence accumulation within a trial and are denoted by arrows. The drift rates for the 36%, 10%, and 0% coherence conditions for both to and away from the IF are denoted as arrows (orange arrows indicate drift rates in the post-muscimol condition and black arrows that indicate drift rates in the pre-muscimol condition). The mean drift rate across coherences is denoted by the longer arrow. As we see in Extended Data Fig. 4a-d, a gain decrease (with the same mean rate of 0) would be evident as an increase in the RT distribution tails for both decisions to and away from the IF (Fig.4a, Extended Data Fig. 4a), a change in the slope of the psychometric function (Fig. 4b, Extended Data Fig. 4b; dots indicate simulated data and curves result from a psychometric curve fit of those simulated data, with black indicating pre-muscimol and orange indicating post-muscimol), and an increase in mean RTs for both correct and error decisions to and away from the IF (Fig. 4c,d, Extended Fig. 4c,d; black indicating pre- and orange indicating post-muscimol). However, we did not observe significant changes in slope of the psychometric curve (Fig. 2f, Fig. 4r,v) nor did we observe consistent mean RT increases for correct choices away from the IF in monkey S (Fig. 4s), making the multiplicative decrease of the drift rate possibility unlikely to explain the post-muscimol data.

The second possibility is that muscimol alters the proportional starting point of the evidence accumulation (*w*) leading to a bias in choosing one or the other option (as a proportion of the boundary parameter, *a*). A change in starting point before evidence accumulation could *also* be considered analogous to the decision criterion in SDT in addition to a change in mean drift rate across coherences but these effects can be differentiated with DDMs due to the different predictions of RT distributions. We simulated a DDM (Fig. 4e-h, Extended Data Fig. 4e-h) with only a decrease in start-point of the evidence accumulation path in proportion to the increasing boundary (pre-muscimol *w*_*1*_ was 0.5 and post-muscimol *w*_*2*_ was 0.25) with the same initial parameters and boundary height change as the other simulations. Note that absolute changes in starting points of evidence accumulation cannot be differentiated from asymmetric changes in evidence boundaries to and away from the IF with behavioral data alone. However changes in the proportional starting point (*w*) *can* be differentiated from symmetric changes in boundaries (*a*)^31^ (see parameter recovery in Supplemental Fig. 2j-m). As we see in Fig.4 e-h and Extended Data Fig. 4e-h of the simulation with a change in start point, the predictions show large changes in the leading edge of the awayIF RT distribution and shorter mean error toIF RTs, along with a decision bias. Looking at the choice performance prediction, a start point change may be a possibility in explaining the effects of muscimol (Fig. 4r,v). However, when we look at the mean error RT’s for both monkeys (Fig. 4t,x), we do not observe a decrease, as suggested by the simulation, but rather an increase, which is suggested by the simulation with a change in mean drift rate across coherences (Fig. 4h, Extended Data Fig. 4h). We also observe a slight shortening of the leading edge of the toIF RT distribution for monkey S, but not for monkey B (Fig. 4q,u). Overall, a change in start point could explain the post-muscimol choice performance data, but explains the RT data less so.

A third possibility is that only the toIF boundary (*aU*) increases (i.e. more evidence is required for a toIF decision after muscimol in the SC) without an absolute start point *(z)* change (Fig.4i-l, Extended Data Fig. 4i-l). Note that an increase in only one decision boundary (perhaps due to inactivation of only one SC) would be reflected as an increase in symmetric boundaries (*a*) and a decrease in proportional start point (*w*) away from the IF with *no changes in mean drift rate across coherences* in the HDDMs and DDMs we used. We simulated an upper boundary change with parameters (*δ*_*1*_ spanned -3 to 3 with steps of 0.5 away with mean *Δ* = 0, *τ* = 0.45 sec, *z* = 0.5, *λ* = 0) with an increase in top boundary from *aU*_*1*_ = 0.7 to *aU*_*2*_ = 1 (which could also be reflected by a symmetric boundary increase of *a*_*1*_ = 1.4 to *a*_*2*_ = 1.7 and proportional start point decrease away from the IF *w*_*1*_ = .50 to *w*_*2*_ = 0.41) shown in Fig. 4i and Extended Data Fig. 4i. A single bound change also predicts a shift in the decision bias (Fig. 4j and Extended Data Fig. 4j), like those seen in the scenario of a proportional start point (*w*) change, the scenario of a change in mean drift rate across coherences (*Δ)*, and also in the actual data (see Fig. 4r,v). For the RT predictions, the single bound change predicts an increase in awayIF RTs for correct and incorrect trials (Fig. 4k,l and Extended Data Fig. 4k,l), which matches the RT change we see in monkey S (Fig. 4s,t) and monkey B (Fig. 4w,x). However, this parameter change cannot explain the magnitude of the psychometric function shift we observed in the data (Fig. 4r,v) with similar changes in simulated and observed mean RTs (Fig. 4s,t,w,x).

The fourth possibility is that muscimol alters or impairs the mean drift rate across coherences, also known as the *zero point of the drift rate*^*31*^ and operationalized as the mean drift rate across positive and negative coherence values (Δ). The mean drift rate across coherences could be considered analogous to the decision criterion in SDT, and alterations in the mean drift rate across coherences would appear as decision biases (in addition to start-point biases, discussed below). We simulated a DDM (Fig. 4m-p, Extended Data Fig. 4m-p) with a change in mean drift rate across coherences (*Δ)*, achieved by an *offset* (additive shift) in drift rates of 1 evidence unit away from the IF (pre-muscimol *δ*_*1k*_ spanned -3 to 3 with steps of 0.5 away with mean *Δ*_*1*_ = 0 and post-muscimol *δ*_*2k*_ spanned -4 to 2 with steps of 0.5 away with mean *Δ*_*2*_ = -1) with the same initial parameters as the other simulations. Note that in this simulation the 0% coherence condition equals both the mean drift rate across coherences (*Δ*_*1*_ = 0 and *Δ*_*2*_ = -1) and the additive offset. A shift in the mean drift rate away from the injected field (IF) in each coherence condition results in a shifted psychometric curve after muscimol injection (Fig. 4n, Extended Data Fig. 4n). The resulting chronometric curve from a shift in drift rates away from the IF is distinct, similar to the shift that we observed in the data (Fig 4r,v). This simulation also predicts an increase in error and correct toIF RTs (Fig. 4o,p, Extended Data Fig. 4o,p), which we observed in both monkeys (Fig 4s,t,w,x), whereas correct awayIF RTs are expected to decrease; an observation made in monkey S but not monkey B. Overall, the mean drift rate change can explain the effects of muscimol in the SC on the choice performance and RT data.

The fifth possibility is that muscimol alters or impairs decision thresholds symmetrically which control the amount of evidence required to reach a decision. We simulated only an increase in symmetric evidence boundaries due to SC inactivation (pre-muscimol *a*_*1*_ was 1.4 and post-muscimol *a*_*2*_ was 1.7) with the same initial parameters as the other simulations (Extended Data Fig. 4q-t). An increase in the symmetric boundary parameter (*a*) always results in an increase of RTs and increases in accuracy whereas a decrease in the symmetric boundary parameter (*a*) always results in a decrease of RTs and decreases in accuracy^31^. Note that a change in sensitivity in SDT could be reflected by a change in symmetric boundary (*a*) or a change in the absolute slope change of drift rates over coherence values. Comparing these symmetric boundary simulation predictions of choice performance and RT to the actual data (Fig. 4q-x), we see a lack of a decision bias prediction, but rather only a slight sensitivity change prediction (Extended Data Fig. 4r). Since we observed a large decision bias in the data, the possibility of a change in symmetric boundaries is unlikely to best explain the effects of muscimol in the SC on the choice performance and RT data.

A sixth possibility is that muscimol inactivation of the SC alters the time for processes within a trial not related to evidence accumulation, ie., non-decision time (*τ*), such as visual encoding of the Glass pattern or saccade execution. We simulated increased non-decision time (pre-muscimol *τ*_*1*_ was 0.45 sec and post-muscimol *τ*_*2*_ was 0.65) with the same initial parameters as the other simulations (Extended Data Fig. 4u-x). A non-decision time change does not predict any changes in SDT because no choice behavior is affected, and therefore is unlikely to explain the post-muscimol data.

These different simulation predictions lay out ways in which SC inactivation can affect RT distributions and decision performance, which we can compare visually with the actual data in Fig 4q-x. However, for a formal analysis comparing predictions of different models, we fit a hierarchical DDM with individual parameters free to vary to see what parameter change best explained the effects of muscimol in the SC.

### Hierarchical drift-diffusion model of decision-making (HDDM)

We estimated parameters of hierarchical drift-diffusion models (HDDMs) fit to pre, post, and recovery data. Hierarchical DDMs often yield better estimates of session parameters due to “shrinkage” towards the mean parameters^41,42^. This is in contrast to fitting a model per session of data which could lead to overfitting and misestimation^43,44^.

The hierarchical drift-diffusion model (HDDM) can be written as a series of statistical relationships between parameters and data. In particular, the HDDM can be written as a set of prior distributions of parameters, statistical parameter relationships, and likelihoods of observed data. Prior distributions provide initial uncertainty about parameters before data are observed. Here, we let only the data influence the shape of the posterior distributions by using wide (less informative) priors for the hierarchical parameters. Notation “∼” refers to a probability distribution of parameters or data. The five probability distributions that were used in the hierarchical model are the Normal distribution (*N*) with mean and standard deviation parameters, the Truncated Normal distribution with truncation between *c* and *d* indicated by *∈ (c, d)*, the Gamma distribution (*Γ*) with rate and shape parameters, the Uniform distribution (*U*) with boundary parameters, and the Wiener likelihood for the drift-diffusion model with parameters: non-decision time (*τ*), symmetric boundary (*a*), proportional start point (*w*), and drift rate (*δ*). Similar to other DDM model fitting procedures^31,45,46^, a lapse process defined by the proportion of trials (*λ*) is found by assuming a mixture between the Wiener likelihood and a uniform distribution over all possible positive and negative RTs less than three seconds (where positive and negative RTs differentiate toIF and awayIF respectively using the Wiener module^37^). This lapse process thus captures a proportion of trials where the monkey did not complete the task and instead made a Bernoulli (i.e. “coin flip”) decision either toIF or awayIF with a random RT between 0 and 3 seconds. The hierarchical mean parameters (*μ*) varied by experimental injection condition (*e*) and monkey (*m*), which affected parameters that varied by experimental session (*s*) with some variance given by hierarchical variance parameters (*σ*). The mean drift rate across coherences (*Δ*) in each experimental session was defined as the mean of the drift rates (*δ*) across positive and negative coherence values (*k*). The overall hierarchical model (HDDM, Supplemental Fig. 1a) of choice and RT data (vector *y*) per trial (*n*) is defined by the following equations:

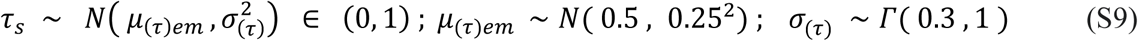

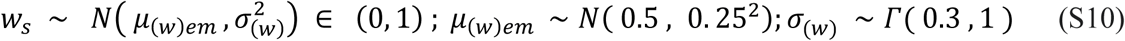

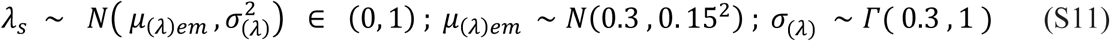

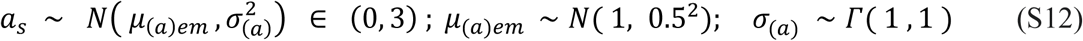

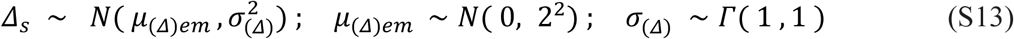

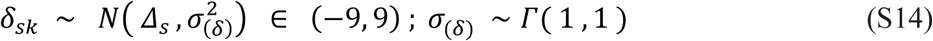

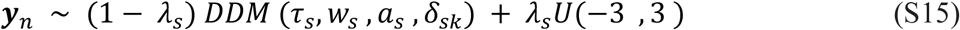

Note that we fit a simple HDDM without trial-to-trial variability in the model parameters, specifically trial-to-trial variability in non-decision time, drift, and initial bias. We assume that some across-trial variability in these parameters exist in the data due to there being intrinsic variability in the stimulus display and the monkeys’ performance. However simple DDM models often estimate the mean DDM parameters well even in the presence of across-trial variability^47^. We therefore simplified the model fitting procedure by excluding across-trial variability parameters, which are often difficult to estimate^48^.

Hierarchical parameter estimates of pre, post, and recovery data are shown in Extended Data Fig 4a-e. Also presented are 2.5th and 97.5th percentiles of the posterior distributions of hierarchical parameters, which provide 95% credible intervals. These credible intervals provide 95% certainty about the parameter estimates given the likelihood of the data and prior distributions. For both monkeys, we see a significant change in the mean drift rate across coherences towards the awayIF decisions in the post-muscimol data after unilateral inactivation of the SC (see Main text, Extended Data Fig 4a). Increases in non-decision time and symmetric bound for monkey B were also observed (Extended Data Fig 4c and d). The changes in non-decision time (*τ*) and symmetric bound (*a*) for monkey S were not very probable (for instance, the 95% credible interval of Post-data includes the estimate of the Pre-data). We did not observe any significant changes in the proportional start point (*w*) (see Main text, Extended Data Fig 4b) nor lapse proportion (*λ*) (Extended Data Fig 4e) for either monkey.

The graphical results of the full HDDM (Fig. 4q,u) were generated by estimating the median posterior samples of each hierarchical parameter (*μ*), the parameter estimates that are shown in Extended Data Fig 4a-e. The graphical DDM for monkey S (Fig. 4q-t) was generated with median posteriors of -0.06 evidence units per second mean drift rate across coherences (*μ*_*Δ*_) pre-muscimol, -0.64 mean drift rate across coherences (*μ*_*Δ*_) post-muscimol, 0.54 proportion of decision evidence as the start-point (*μ*_*w*_) pre-muscimol, 0.49 proportion of decision evidence as the start-point (*μ*_*w*_) post-muscimol, 1.50 evidence units boundary (*μ*_*a*_) pre-muscimol, 1.55 evidence units boundary (*μ*_*a*_) post-muscimol, 408 ms non-decision time (*μ*_*τ*_) pre-muscimol, 433 ms non-decision time (*μ*_*τ*_) post-muscimol. The median posteriors of 3% lapse rate (*μ_λ_*) pre-muscimol and 4% lapse rate (*μ_λ_*) post-muscimol for monkey S are not shown in the graphical results (Fig. 4q). The graphical DDM for monkey B was generated with mean posteriors of 0.09 mean drift rate across coherences (*μ*_*Δ*_) pre-muscimol, -0.85 mean drift rate across coherences (*μ*_*Δ*_) post-muscimol, 0.52 proportion of decision evidence as the start-point (*μ*_*w*_) pre-muscimol, 0.55 proportion of decision evidence as the start-point (*μ*_*w*_) post-muscimol, 1.33 evidence units boundary (*μ*_*a*_) pre-muscimol, 1.55 evidence units boundary (*μ*_*a*_) post-muscimol, 543 ms non-decision time (*μ*_*τ*_) pre-muscimol, 597 ms non-decision time (*μ*_*τ*_) post-muscimol. The median posteriors of 5% lapse rate pre-muscimol and 5% lapse rate post-muscimol. We also estimated the median posterior samples of drift rates for each monkey in each session by combining the posterior distributions across sessions. The estimates were for the -36%, -10%, 0%, 10%, and 36% were used to plot drift rate estimates as arrows in the graphical representation (Fig. 4q,u) and were the following: (−1.82, -0.86, -0.16, 0.71, 1.96) pre-muscimol in monkey S, (−1.89, -1.29, -0.90, -0.02, 1.3) post-muscimol in monkey S, (−2.74, -1.13, 0.20, 1.71, 3.29) pre-muscimol in monkey B, and (−3.20, -1.87, -0.87, 0.02, 2.16) post-muscimol in monkey B.

Given that there are multiple parameter changes from pre-muscimol to post-muscimol, we tested which parameter best described the observed behavioral changes in choice and RT distributions after muscimol injection in the SC by fitting the HDDMs that fixed all parameters across injection conditions besides the parameter of interest. This resulted in four additional versions of the hierarchical DDM that were fit to the data: HDDM with the drift rate mean across coherences free to vary (HDDM-*Δ*), HDDM with the start point free to vary (HDDM-*w*), HDDM with non-decision time free to vary (HDDM*-τ*), and HDDM with the start point and bound free to vary (HDDM-*a,w*). In each new model, hierarchical parameters (*µ*), besides hierarchical parameters of the variable of interest (*Δ* or *w* or *τ*), only varied by monkey (*m*) and *not* by injection condition (*e*). Hierarchical parameters of the variable of interest could vary by both monkey (*m*) and injection condition (*e*). Sessions (*s*) were collapsed across the pre- and post-muscimol and recovery sessions for each separate injection into a parameter “day” (*d*), where one “day” collapsed across three sessions of data that constituted two sessions (a pre-session occurring immediately preceding a post-muscimol or saline session) and a recovery session within ∼24 hours. Parameters besides the variable of interest could vary by day (*d*), while the parameter of interest varied by session (*s*). Thus, each parameter besides the parameter of interest could only explain variance in the data due to the monkey (B or S) and across unique “days”. Only the parameter of interest could explain variance in the data due to the effect of an injection changing across injection types (*e*) and separate sessions (*s*). Similar non-informative prior distributions were used as in the full HDDM. The overall hierarchical model with mean drift rate across coherences (*Δ*) as the variable of interest (HDDM-*Δ*, Supplemental Fig. 1b) of choice and RT data (vector *y*) per trial (*n*) is defined by the following equations:

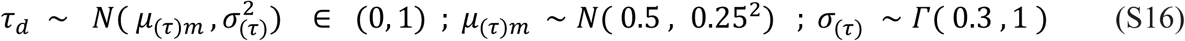

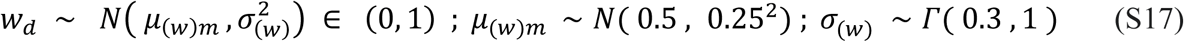

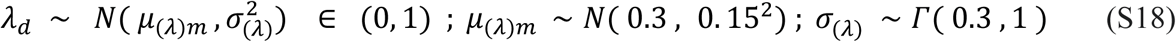

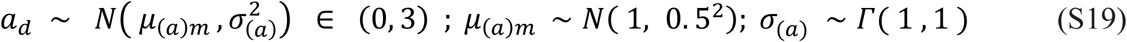

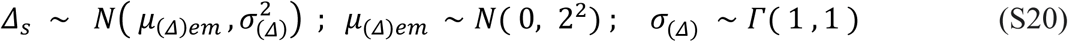

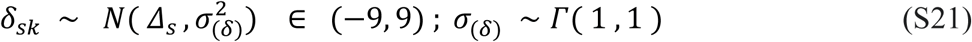

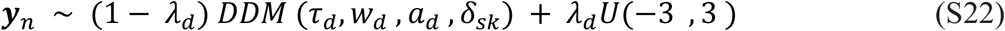

As in the full HDDM, in a hierarchical model with start point (*w*) as the variable of interest (HDDM-*w*, Supplemental Fig. 1c), the start point of evidence accumulation (*w*) was assumed to be fixed across coherence values before evidence accumulation in a trial. The model is defined by the following equations:

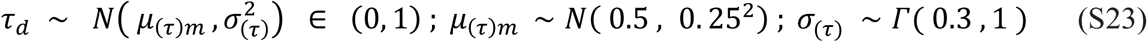

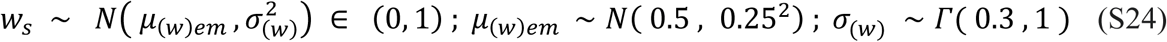

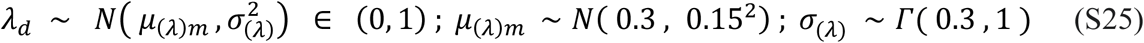

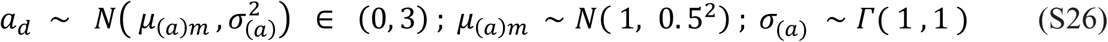

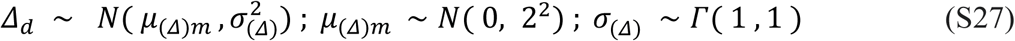

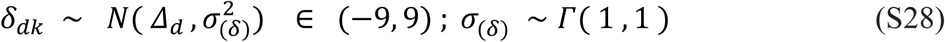

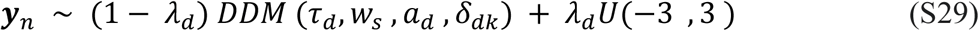

Unlike in the full HDDM, non-decision time (*ω*) was allowed to be variable across coherence values (*k*) with a hierarchical non-decision time (*τ*) as the variable of interest (HDDM*-τ*, Supplemental Fig. 1d). The model is defined by the following equations:

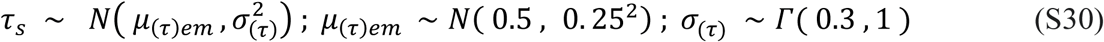

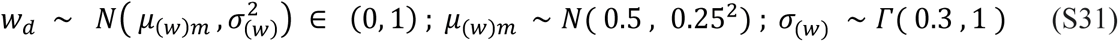

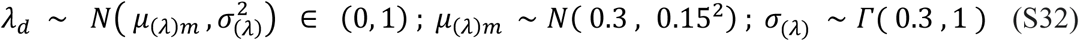

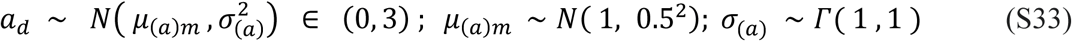

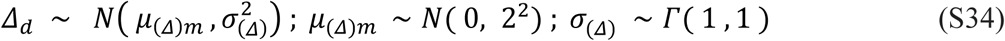

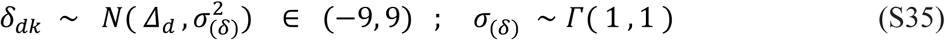

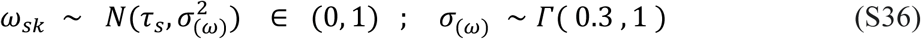

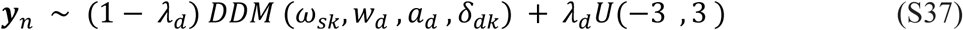

To answer the question whether *any* boundary change (including a symmetric boundary or either single boundary change) could explain the effects of unilateral inactivation of the SC with muscimol, we fit a model with only the symmetric boundary (*a*) and start point (*w*) free to vary across experimental conditions (*e*) and sessions (*s*). This final model is defined by the following priors and likelihood equations:

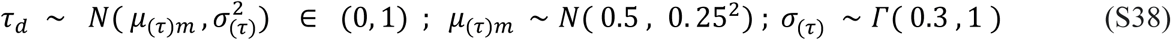

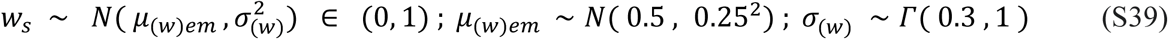

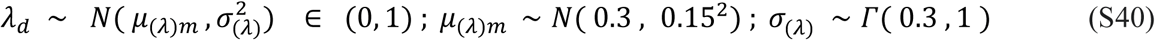

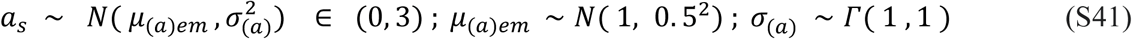

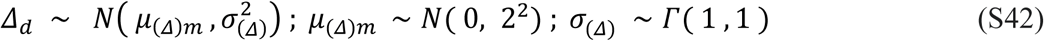

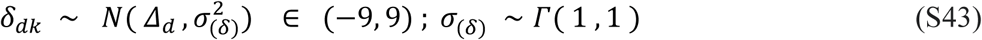

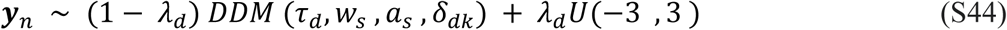

When comparing the in-sample prediction of all five model variants, the full HDDM explains the change in decision bias the best in both monkeys (98.3% of the data for monkey S, 99.3% for monkey B) as shown in the predicted psychometric functions (Extended Data Fig. 6; Extended Data Table 4). The HDDM-*Δ* captures the change in decision bias almost as well as the full HDDM in both monkeys (97.6% for monkey S, 98.3% for monkey B), followed by HDDM-*a,w* (96.4% for monkey S, 94.0% for monkey B), then HDDM-*w* (94.5% for monkey S, 93.5% for monkey B), and then HDDM-*τ* (89.9% for monkey S, 91.4% for monkey B). The full HDDM also explains the RT distribution best for both monkeys (Extended Data Table 4) as seen by the visual comparisons of the predictions of the HDDM and the actual data (Extended Data Fig. 6). While HDDM-*τ* fails to capture any decision bias from pre to post data, it does capture RT data almost as well as the full model (Extended Data Fig. 6; Extended Data Table 4). Overall the full HDDM best predicts the choice and the RT data and the results of this model are discussed in the Main text.

**Supplemental Fig. 1.**
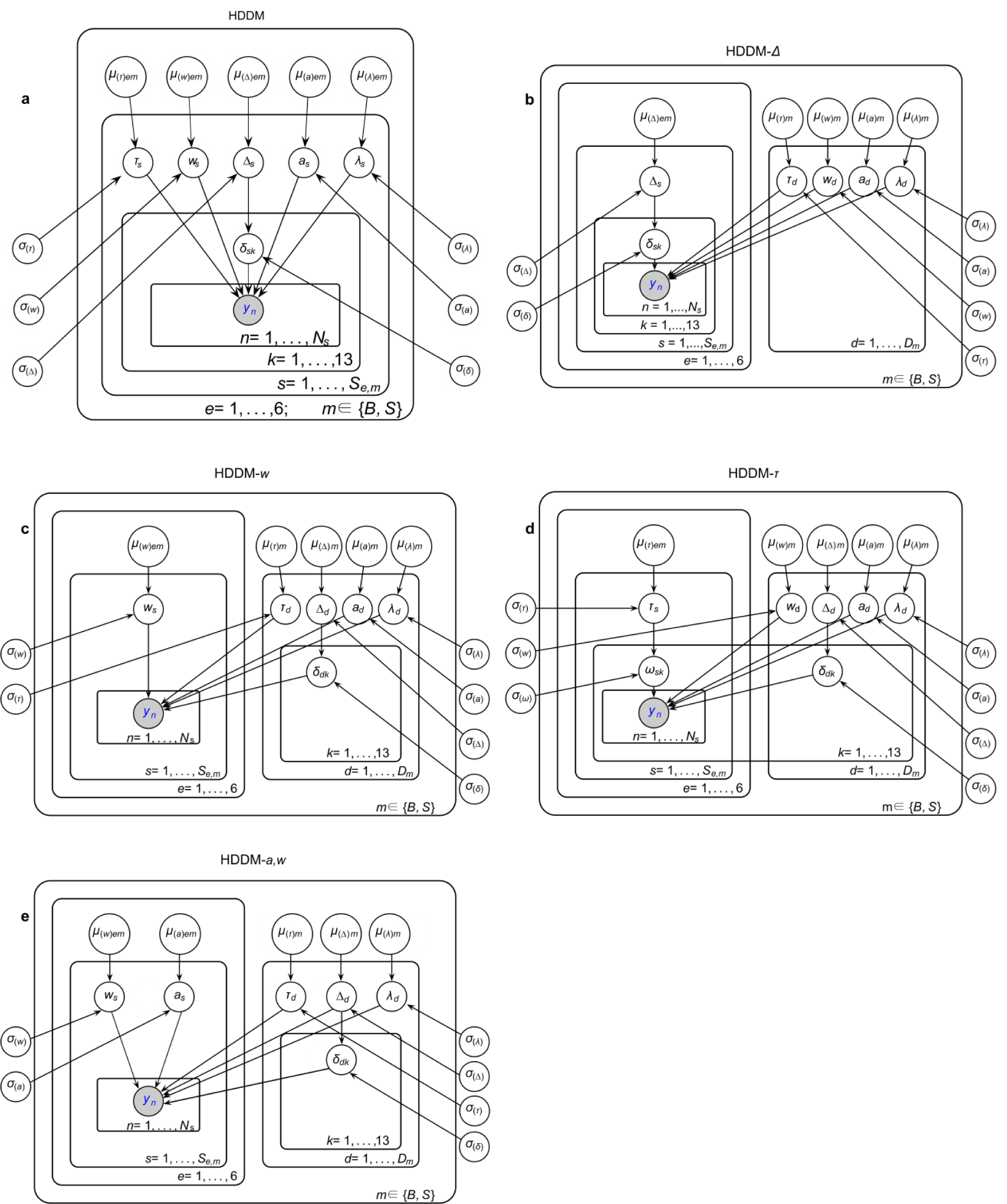
Graphical model descriptions. (Associated with Fig. 4 of the main text) Hierarchical models represented by the graph structure^45^ of **a** full HDDM **b** HDDM-*Δ* **c** HDDM-*w* **d** HDDM-*τ* **e** HDDM-*a,w*, given by the prior and likelihood equations in the Supplementary Materials (S9-44). Arrows represent dependencies of variables on hierarchical variables, including hierarchical means (*μ*) and hierarchical variance parameters (*σ*). Hierarchical means vary by either both monkey (*m*) and injection condition (*e*) or just monkey (*m*). Other variables vary by session (*s*), day (*d*), coherence condition (*k*). The observed joint choice-RT data (***y***) per trial (*n*) is described by a Wiener process depending upon variables in the lowest level of the hierarchy.

To better understand the mechanism of unilateral inactivation of the SC on evidence accumulation, we fit one final HDDM with the same parameters and priors as the full HDDM but with an enforced linear equation that describes each drift rate (*δ*) per experimental session (*s*) and coherence *(k*) as the linear comparison of two accumulators. One can interpret this model as the addition of a gain element that impacts how evidence is accumulated in one of two accumulators ^50,51^. To achieve model identifiability, we assumed that the injected SC accumulated a probability for an IF decision with probability (*θ*) that varied with coherence *(k*) with the other SC accumulating with probability (1 - *θ*). The injected SC was also influenced by gain effects (*G*_*(toIF)*_) and the unaffected SC was influenced by gain effects (*G*_*(awayIF)*_) that both varied by injection condition (*e*). For model identifiability, we could only gauge the relative influence of injection conditions on the gain effects, and thus *G*_*(awayIF)*_ was set to 1 in each injection condition and *G*_*(toIF)*_ was set to 1 in the Pre-muscimol injection condition. We explored the amount of gain on toIF decisions (*G*_*(toIF)*_) in the other five experimental conditions: Post-muscimol, Recovery from muscimol, Pre-saline, Post-saline, and Recovery from saline with Bayes Factors (see Methods). Finally, the linear accumulator comparison was scaled by a parameter in seconds (*R*) that varied per monkey (*m*). Thus, this final model (HDDM-*G*) is fully described by Equations S9 through S12 as well as the following equations:

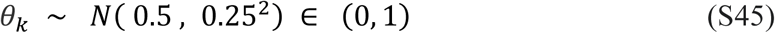

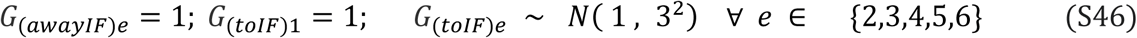

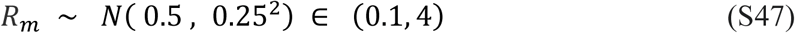

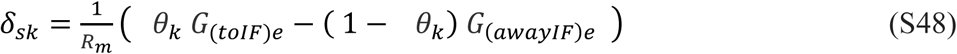

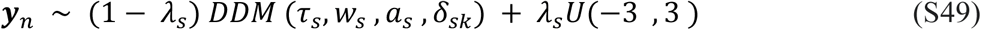

All HDDMs were fit for 52,000 original samples in each of six Markov chains for each parameter. After removing the first 2000 samples as a “warm-up” and then keeping only every 10th sample, i.e. using a “thinning” parameter of 10, this resulted in 5,000 posterior samples in each Markov chain for 5,000 * 6 = 30,000 samples from the estimated posterior distributions for each parameter. To assess model convergence, the Gelman-Rubin statistic and the number of effective samples were calculated^44^. The Gelman-Rubin statistic assesses the convergence of MCMC samplers by comparing the between-chain variance to the within-chain variance of each parameter, with Gelman-Rubin statistic > 1.1 thought to be a necessity for convergence. The “effective number of samples” equation scales the total sample number for each parameter posterior by autocorrelation in the chains in order to estimate an independent number of samples^44^. We also implemented the recommendation by Gelman et al.^44^ (see footnote in the 3rd Edition on page 283) to split the chains in half before calculating the Gelman-Rubin statistic in order to account for non-stationary chains. Larger effective numbers of samples for each parameter in the model are better. The chains for parameters with the largest Gelman-Rubin statistics and smallest effective number of samples were also visually inspected to ensure convergence. Model HDDM converged with Gelman-Rubin statistics of <= 1.03 and minimum number of effective samples of >=222 for all parameters. Model HDDM-*Δ* converged with Gelman-Rubin statistics of <= 1.01 and a number of effective samples of >=882 for all parameters, and model HDDM-*w* converged with Gelman-Rubin statistics of <= 1.03 and a number of effective samples of >=31 for all parameters. Model HDDM-*τ* had some non-decision time variables reach multiple-peaked posterior distributions, and so did not converge with Gelman-Rubin statistics of <= 1.12 and a number of effective samples of >=37 for all parameters. The model HDDM-*a,w* converged with Gelman-Rubin statistics of <= 1.06 and a number of effective samples of >=22 for all parameters. Model HDDM-*G* converged with Gelman-Rubin statistics of <= 1.02 and a number of effective samples of >=337 for all parameters. The goodness of fit measure, *R*^*2*^_*pred*_ for these sets of parameter estimates for HDDM to the data are shown in Extended Data Table 4.

### Non-hierarchical drift-diffusion model of decision-making (DDM)

We also fit the non-hierarchical DDM, to the pre-muscimol and post-muscimol data using quantile maximum probability estimation (QMPE)^49^ in the ChaRTr package^38^, along with the HDDM to ensure that our parameter estimations were robust against different modeling methods. Model fitting of the non-hierarchical DDM was performed on the RT task data from monkey S and monkey B (seven for monkey S and two for monkey B), separately for each monkey for pre- and post-muscimol data, and pooled across sessions. This model fit to the data with nine RT quantiles that summarize the RT distribution of each of the 11 trial conditions (24%, 17% 10%, 5%, 3%, 0%) for toIF and awayIF decisions with the 0% collapsed for to and awayIF decisions. The 36% conditions in the toIF and awayIF data were excluded for both monkeys because there were not enough data for fitting error RT quantiles. For all the fits using the models in the ChaRTr package, five iterations were fit for each dataset with different random seeds to avoid getting stuck in local minima when optimizing the goodness of fit measure, the QMP statistic. The best out of the five fits (according to AIC/BIC scores calculated from the QMP statistic) was used for model comparisons and reported for parameter estimation (Extended Data Table 4 and Extended Data Fig. 5).

The main differences between the HDDM and the non-hierarchical DDM are that 1) they employ a different modeling method (QMPE for non-hierarchical DDM and Bayesian Estimation for HDDM), 2) the HDDM has a hierarchical structure that allows for the parameters of individual experimental sessions to be pushed towards the mean parameters of all the sessions in an effort to better estimate parameters, and 3) the inclusion of a lapse rate in the HDDM. For our non-hierarchical DDM, the drift rate for each condition, start point (*w*), non-decision time (*τ*), and the symmetric boundary (*a*) were free parameters that were estimated for pre- and post-muscimol data.

The parameter estimates of the DDM from fitting pre and post sessions showed the same relative changes from pre- to post-muscimol in all the parameters for both monkeys that we also saw using the HDDM parameter estimates, including the same change in the mean drift rate across coherences (Extended Data Fig 4f), indicating that our parameter estimation results are robust against different modeling methods.

### Urgency-gating Models (UGMs)

Urgency-gating Models (UGMs) are another class of accumulation to bound model where the sensory evidence is both 1) low-pass filtered to prioritize more recent evidence and 2) multiplied by a linearly growing urgency signal during a decision^13,14^. We found evidence that the UGM may also reasonably explain the behavior of monkey B in the pre-muscimol sessions, whereas data from monkey S were better explained by DDMs (Extended Data Table 4), although it is impossible to distinguish DDMs from UGMs using tasks with constant evidence during a trial, such as the Glass Pattern decision task^13,14^. Nevertheless, under the assumption that the monkeys’ behavior could be explained by the UGM, and given our finding that a change in the mean drift rate across coherences (*Δ*) best explained the effect of the muscimol when assuming a DDM (Extended Data Table 4), we tested whether unilateral SC inactivation could be explained by a change in mean drift rate across coherences or a change in the urgency signal of a UGM. To assess this, we fitted UGMs to the data from monkey S and monkey B by keeping all parameters fixed post-muscimol from their pre-muscimol values, but with either a free-to-vary urgency slope parameter (UGM-*m*) or a free drift rate across coherence values (UGM-*δ*) to see whether a change in evidence accumulation or a change in urgency signal better explained the data.

This UGM process that we used for our model is defined by the following equations modified from the ChaRTr package:

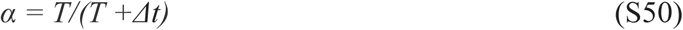

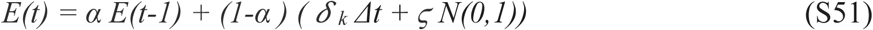

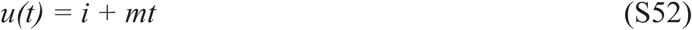

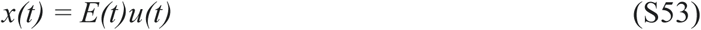

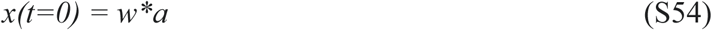

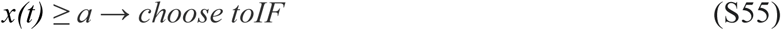

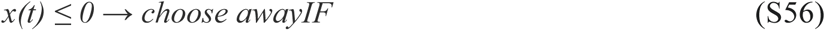

Evidence during a trial, *E(t)*, is low pass filtered with a weight for previous evidence (*α*) that is controlled by the filter constant set to *T =* 100 ms and a weight for incoming evidence (1-*α*) that is influenced by the drift rate (*δ*) for each condition (*k*), the time step (*Δt* = 1 ms), and the noise constant (*ϛ*), set to 100 evidence units. The current state of evidence, *E(t)*, is multiplied by the linear urgency signal *u(t)* with a slope (*m*) and an intercept (*i*), fixed to 0, to make the decision variable, *x(t)*.

The slope of the linearly growing urgency signal (*m*) in the UGM was free to vary to assess whether the effects of muscimol were best explained by a change in the urgency signal. The symmetric boundary (*a*) was fixed to estimate the urgency slope (*m*) as a free parameter because parameter recovery results of the UGM with both a free boundary and a free urgency slope suggested that there was an overlapping role of the bound height and the urgency slope in explaining the choice and reaction time data (Supplemental Fig. 2a-e). Because we were only interested in assessing whether the effects of muscimol in the SC were best explained by a change in the urgency signal or the drift rate, we fixed the symmetric boundary in order to accurately estimate the change in urgency slope from pre- to post-muscimol. For fitting the UGM with both the pre- and post-muscimol data, the boundary value was fixed at the value of the parameter estimate of the boundary from fitting the standard UGM with the only the boundary (*a*), drift rates (*δ* _*k*_), non-decision time (*τ*), and start point (*w*) allowed to vary in the pre-muscimol data. Therefore, in our full UGM, the drift rate for each condition (*δ* _*k*_), non-decision time (*τ*), start point (*w*), and the slope of the urgency signal (*m*) were free parameters that were estimated for pre- and post-muscimol data for both monkeys.

**Supplemental Fig. 2.**
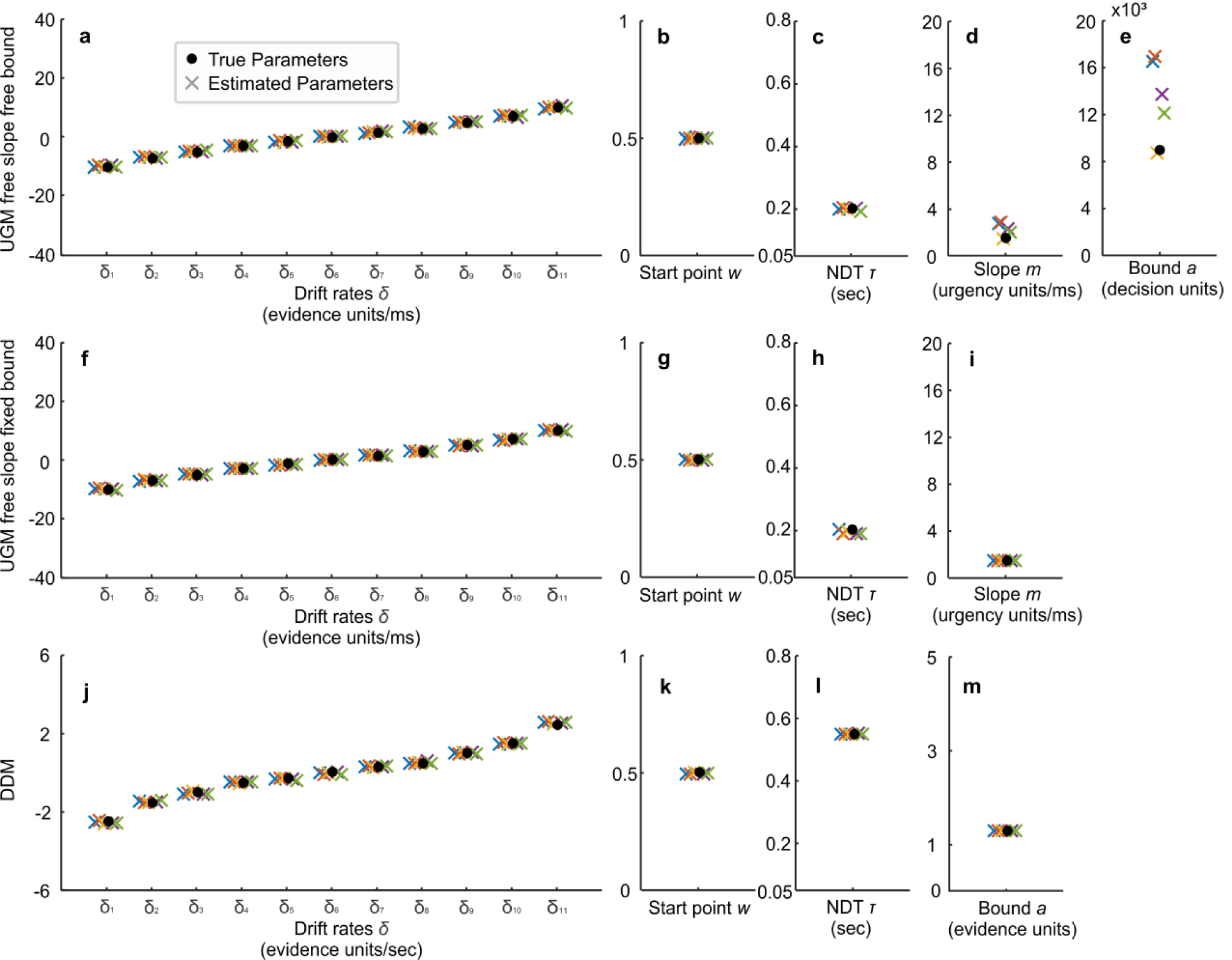
Parameter Recovery for UGM and DDM. The set of parameter estimates of each of the five iterations of fits for parameter recovery (denoted as x) are shown along with the known parameters used to simulate the data (denoted as black circles). The y-axis scale for each parameter was determined by the lower restraint and the upper restraint when estimating the parameter during fitting. **a-e** The parameter recovery estimates for the UGM with free proportional start point, free drift rates, free non-decision time (NDT), free urgency slope, and a free bound. Visually, the parameter recovery estimates of all 5 iterations of the drift rate, proportional start point, and non-decision time match the true parameter estimate. However, parameter estimates from four out of the five fits of the urgency slope and the bound are overestimated. The magnitude of the overestimation of the urgency slope and the bound even seem correlated, where the fits with a higher slope also have a higher bound. Because we were unable to recover the slope when the bound was free reliably, and because we want to assess whether a change in urgency slope best explained the effect of muscimol in the SC, we used a version of the UGM where the slope, drift rate, proportional start point, and non-decision time were free, but the bound was fixed. **f-i** To confirm that the UGM with the free slope but fixed bound (proportional free start point, free drift rates, free non-decision time, free slope, fixed bound) was able to estimate parameters accurately, we show the parameter recovery estimates for this UGM version. All parameter recovery estimates of all five fits recover the true parameter for this model. We defined this as our “full UGM” and used for modeling the actual data to recover slope estimates. **j-m** We also made sure that the non-hierarchical DDM can recover parameters accurately with free proportional start point, free drift, free bound, and free non-decision time. All parameters from all of the 5 fits match the true parameters.

For fitting the UGM for parameter estimation of pre- and post-muscimol data and also for fitting UGM variants with only either the drift rate or the urgency slope free, we used the same pre- and post-musimol data that was pooled across sessions and modeling method described in the non-hierarchical DDM section, using the ChaRTr package^38^. To test the robustness of this procedure of fitting pooled data, we also fit all the pre-muscimol sessions individually across experimental sessions, with only 9 conditions: 17%, 10%, 5%, 3% each for toIF and awayIF, and 0% condition collapsed across toIF and awayIF. The 24% and 36% conditions were excluded because there were not enough error trials in each individual session for either monkey. We also used the results of fitting individual pre-muscimol session data, for model comparison between the HDDM, DDM, and UGM for each monkey (model fits denoted with ** in Extended Data Table 4). The parameter estimates for the DDM and the UGM for the pooled and individual data were similar (Supplemental Fig. 3).

**Supplemental Fig. 3.**
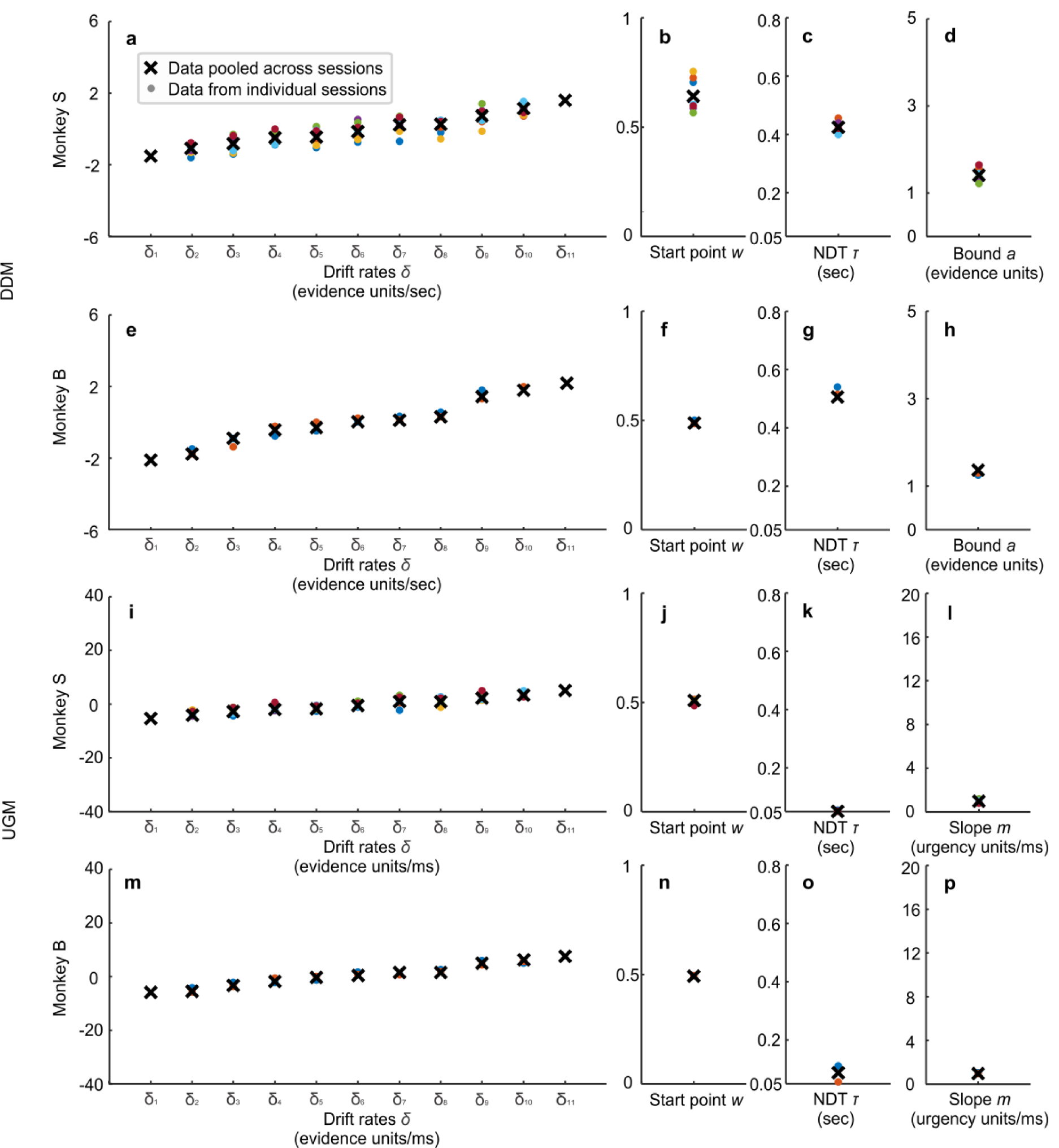
Parameter estimates of fits using pre-muscimol data from individual data versus pooled data. **a-d** DDM parameter estimates fit to pre-muscimol data that were pooled across all muscimol injections (black x) are shown in comparison to the DDM parameter estimates fit to the pre-muscimol data from individual muscimol injection sessions for monkey S (n = 9, circles). **e-h** Same as in a-d for monkey B (n = 2, circles). **i-l** Same as in a-d for the UGM for monkey S (n = 9, circles). **m-p** Same as in i-l for monkey B (n = 2, circles). For both monkeys and for both models, although there is some variability in parameter estimates between injection sessions, especially in DDM parameter estimates for monkey S, the parameter estimates obtained using pooled data for fitting the models summarize the parameters from all the individual sessions reasonably well.

The parameter estimates of the UGM from fitting pre- and post-muscimol sessions show the relative changes from pre- to post-muscimol for the mean drift rate across coherences as seen with the HDDM and DDM, along with changes in urgency slope for both monkeys, slightly for monkey S and more so for monkey B (Extended Data Fig. 5k,l). However, when fitting the UGM with only the slope free to vary, the model failed to capture the post-muscimol decision bias data (Extended Data Table 4 and Extended Data Fig 6b and d in UGM-*m* row). In contrast, the UGM with the drift rates free to vary captured the choice data almost as well as the full UGM (Extended Data Table 4 and Extended data Fig 6b and d in UGM-*δ* row), indicating that the change in mean drift rate across coherences best explains the effect of muscimol on decision bias, and confirming the results obtained using the HDDM and DDM.

### Model Comparison

To compare across models, we found the amount of variance explained for in-sample and out-of sample prediction of choice-RT data (Extended Data Table 4). In-sample prediction refers to how well each model performed at predicting proportion of choices and RT quantiles of the data that were used to fit model parameters. We used 80% of the in-session trials (randomly picked with a uniform distribution), to fit model parameters and determine in-sample prediction statistics. Out-of-sample predictions were obtained using the 20% of the data that was not used to fit the data. Bayesian Information Criteria (BIC) and Akaike Information Criteria (AIC) were also calculated for the DDM and UGM models in ChaRTr, which are measures of in-sample prediction collapsed across data types (choice performance and RT) that include penalties for model complexity. BIC has been shown to more often favor models that match the ground truth while AIC has been shown to more often favor models that will be more predictive of new data^38^.

*R*^*2*^_*pred*_ measures the amount of variance in the data (either in-sample or out-of-sample), explained by the prediction of the model. Obtaining estimates of variance explained for RT statistics (mean, 25th percentiles, median, and 75th percentiles) and proportions of choice are useful to understand how one model performs better than another model. *R*^*2*^_*pred*_ ranges from -∞ to 1 and can be multiplied by 100 for an estimate of -∞ to 100% percentage of variance explained by the data statistic (*Q*) by prediction. Note that values can be negative indicating that the amount of variance in prediction is greater than the variance of the statistic (Extended Data Table 4). However, the relative values from one model to the next are still informative about the improvement in prediction. *R*^*2*^_*pred*_ is defined as one minus the mean squared error of prediction of a certain statistic (*Q*) scaled by the estimated variance of a statistic using a number of observations *J* equal to the total number of coherence conditions (*K*) across experimental sessions (*S*) for a total *J = K*S*.

For calculating the *R*^*2*^_*pred*_ for the non-hierarchical DDM and UGM that were fit to individual pre-muscimol sessions for model comparison between the HDDM, DDM, and UGM, we used the individual session’s predictions (with the 24% and 36% conditions excluded, see UGMs section) in calculating the mean squared error of prediction for *R*^*2*^_*pred*_, while the *R*^*2*^_*pred*_ of the HDDM was also calculated from fitting the exact pre-muscimol data for each monkey without the 36% and 24% coherence conditions (model fits denoted with ** in Extended Data Table 4). For all other *R*^*2*^_*pred*_ for the non-hierarchical DDM and the UGMs (i.e. DDM, UGM, UGM-*δ*, UGM-*m*), we found the difference between the predictions from those pooled fits to each individual session’s actual data to calculate the error of prediction.

### Parameter recovery

We performed parameter recovery to make sure that QMPE would return accurate estimates of free-to-vary UGM and DDM parameters. For our process of parameter recovery, we first simulated choice and RT data for each model, using parameter values that were approximated from previous fits of our pre-muscimol data for monkey B using the ChaRTr original DDM and UGM fits of data. We then fit our modified UGM and DDM that we use in our study to those simulated data to discover whether the parameter estimates matched the true parameters. All five sets of parameter estimates from the five fits demonstrated that the models used (DDM and UGM with fixed bound and free slope) can recover parameters accurately. We found that the parameter estimates closely matched the known parameters for the DDM and the UGM (Supplemental Fig. 2).

## Main references

1 Glass, L. The Moire effect from random dots. Nature 223, 578–580, doi: 10.1038/223578a0 (1969).

2 Smith, M. A., Kohn, A. & Movshon, J. A. Glass pattern responses in macaque V2 neurons. J Vis 7, 5, doi: 10.1167/7.3.5 (2007).

3 Hikosaka, O. & Wurtz, R. H. The basal ganglia. Rev Oculomot Res 3, 257–281 (1989).

4 Lee, C., Rohrer, W. H. & Sparks, D. L. Population coding of saccadic eye movements by neurons in the superior colliculus. Nature 332, 357–360, doi: 10.1038/332357a0 (1988).

5 McPeek, R. M. & Keller, E. L. Saccade target selection in the superior colliculus during a visual search task. J Neurophysiol 88, 2019–2034 (2002).

6 Lovejoy, L. P. & Krauzlis, R. J. Inactivation of primate superior colliculus impairs covert selection of signals for perceptual judgments. Nat Neurosci 13, 261–266, doi: 10.1038/nn.2470 (2010).

7 Basso, M. A. & May, P. J. Circuits for action and cognition: A view from the superior colliculus. Annual Review of Vision Science 3, 197–226, doi: 10.1146/annurev-vision-102016-061234 (2017).

8 Krauzlis, R. J., Lovejoy, L. P. & Zénon, A. Superior colliculus and visual spatial attention.Annu Rev Neurosci 36, 165–182, doi: 10.1146/annurev-neuro-062012-170249 (2013).

9 Kim, B. & Basso, M. A. A probabilistic strategy for understanding action selection. J Neurosci 30, 2340–2355, doi: 10.1523/jneurosci.1730-09.2010 (2010).

10 Herman, J. P., Katz, L. N. & Krauzlis, R. J. Midbrain activity can explain perceptual decisions during an attention task. Nat Neurosci 21, 1651–1655, doi: 10.1038/s41593-018-0271-5 (2018).

11 Nummela, S. U. & Krauzlis, R. J. Inactivation of primate superior colliculus biases target choice for smooth pursuit, saccades, and button press responses. J Neurophysiol 104, 1538–1548, doi: 10.1152/jn.00406.2010 (2010).

12 Sedaghat-Nejad, E., Herzfeld, D. J. & Shadmehr, R. Reward prediction error modulates saccade vigor. J Neurosci 39, 5010–5017, doi: 10.1523/jneurosci.0432-19.2019 (2019).

13 Cisek, P., Puskas, G. A. & El-Murr, S. Decisions in changing conditions: the urgency-gating model. J Neurosci 29, 11560–11571, doi: 10.1523/jneurosci.1844-09.2009 (2009).

14 Evans, N. J., Hawkins, G. E., Boehm, U., Wagenmakers, E. J. & Brown, S. D. The computations that support simple decision-making: A comparison between the diffusion and urgency-gating models.Sci Rep 7, 16433, doi: 10.1038/s41598-017-16694-7 (2017).

15 Ratcliff, R. & McKoon, G. The diffusion decision model: theory and data for two-choice decision tasks. Neural Comput 20, 873–922, doi: 10.1162/neco.2008.12-06-420 (2008).

16 Felsen, G. & Mainen, Z. F. Neural substrates of sensory-guided locomotor decisions in the rat superior colliculus. Neuron 60, 137–148, doi: 10.1016/j.neuron.2008.09.019 (2008).

17 Crapse, T. B., Lau, H. & Basso, M. A. A role for the superior colliculus in decision criteria. Neuron 97, 181-194.e186, doi: 10.1016/j.neuron.2017.12.006 (2018).

18 Zhou, Y. & Freedman, D. J. Posterior parietal cortex plays a causal role in perceptual and categorical decisions. Science 365, 180–185 (2019).

19 Gandhi, N. J. & Katnani, H. A. Motor functions of the superior colliculus. Annu Rev Neurosci 34, 205–231, doi: 10.1146/annurev-neuro-061010-113728 (2011).

20 Katz, L. N., Yates, J. L., Pillow, J. W. & Huk, A. C. Dissociated functional significance of decision-related activity in the primate dorsal stream. Nature 535, 285–288, doi: 10.1038/nature18617 (2016).

21 Carandini, M. & Churchland, A. K. Probing perceptual decisions in rodents. Nat Neurosci 16, 824–831, doi: 10.1038/nn.3410 (2013).

22 Hanks, T. D. et al. Distinct relationships of parietal and prefrontal cortices to evidence accumulation. Nature 520, 220–223, doi: 10.1038/nature14066 (2015).

23 Brody, C. D. & Hanks, T. D. Neural underpinnings of the evidence accumulator. Curr Opin Neurobiol 37, 149–157, doi: 10.1016/j.conb.2016.01.003 (2016).

24 Ditterich, J. Evidence for time-variant decision making. European Journal of Neuroscience 24, 3628–3641, doi: 10.1111/j.1460-9568.2006.05221.x. (2006).

25 Ratcliff, R. et al. Dual diffusion model for single cell recording data from the superior colliculus in a brightness discrimination task. J Neurophysiol 97,(2) 1756-1774. doi.org/10.1152/jn.00393.(2006).

## Methods references

24 Judge, S., Richmond, B. & Chu, F. Implantation of magnetic search coils for measurement of eye position: an improved method. Vision Research 20, 535–538, doi: 10.1016/0042-6989(80)90128-5 (1980).

25 Hays, A. V., Richmond, B. J. & Optican, L. M. Unix-based multiple-process system, for real-time data and control. (1982).

26 Fuchs, A. F. & Robinson, D. A. A method for measuring horizontal and vertical eye movement chronically in the monkey. J. Appl. Physiol. 21, 1068–1070, doi: 10.1152/jappl.1966.21.3.1068 (1966).

27 Crist, C. F., Yamasaki, D. S. G., Komatsu, H. & Wurtz, R. H. A grid system and a microsyringe for single cell recording. J. Neurosci. Methods 26, 117–122, doi: 10.1016/0165-0270(88)90160-4 (1988).

28 Ferster, C. B. & Skinner, B. F. Schedules of reinforcement. (Appleton-Century-Crofts, 1957).

29 Ditterich, J. Evidence for time-variant decision making. European Journal of Neuroscience 24, 3628–3641, doi: 10.1111/j.1460-9568.2006.05221.x. (2006).

30 Macmillan, N. A. & Creelman, C. D. Detection theory: A user’s guide. 2 edn, (Lawerence Erlbaum Associates, Inc., 2004).

31 Ratcliff, R., Smith, P. L., Brown, S. D. & McKoon, G. Diffusion decision model: Current issues and history. Trends in Cognitive Science 20, 260–281, doi: 10.1016/j.tics.2016.01.007 (2016).

32 O’Connell, R. G., Shadlen, M. N., Wong-Lin, K. & Kelly, S. P. Bridging neural and computational viewpoints on perceptual decision-making. Trends Neurosci 41, 838–852, doi: 10.1016/j.tins.2018.06.005 (2018).

33 Voss, A., Rothermund, K. & Voss, J. Interpreting the parameters of the diffusion model: An empirical validation. Memory & Cognition 32, 1206–1220, doi: 10.3758/bf03196893 (2004).

34 Plummer, M. in 3rd International Workshop on Distributed Statistical Computing (eds Kurt Hornik, Friedrich Leisch, & Achim Zeileis) 10 (Vienna, Austria, 2003).

35 Prins, N. & Kingdom, F. A. A. Applying the model-comparison approach to test specific research hypotheses in psychophysical research ssing the Palamedes toolbox. Front Psychol 9, 1250, doi: 10.3389/fpsyg.2018.01250 (2018).

36 Miasko, T. PyJAGS: The Python Interface to JAGS, <https://github.com/tmiasko/pyjags> (2017).

37 Wabersich, D. & Vandekerckhove, J. Extending JAGS: a tutorial on adding custom distributions to JAGS (with a diffusion model example). Behav Res Methods 46, 15–28, doi: 10.3758/s13428-013-0369-3 (2014).

38 Chandrasekaran, C. & Hawkins, G. E. ChaRTr: An R toolbox for modeling choices and response times in decision-making tasks. J Neurosci Methods 328, 108432, doi: 10.1016/j.jneumeth.2019.108432 (2019).

39 Shen, Y. & Richards, V. M. A maximum-likelihood procedure for estimating psychometric functions: thresholds, slopes, and lapses of attention. J Acoust Soc Am 132, 957–967, doi: 10.1121/1.4733540 (2012).

40 Gold, J. I. & Ding, L. How mechanisms of perceptual decision-making affect the psychometric function. Prog Neurobiol 103, 98–114, doi: 10.1016/j.pneurobio.2012.05.008 (2013).

## Supplementary methods references

41 Ratcliff, R. & Childers, R. Individual differences and fitting methods for the two-choice diffusion model of decision making. Decision (Wash D C) 2015, doi: 10.1037/dec0000030 (2015).

42 Vandekerckhove, J., Tuerlinckx, F. & Lee, M. D. Hierarchical diffusion models for two-choice response times. Psychol Methods 16, 44–62, doi: 10.1037/a0021765 (2011).

43 Gelman, A. et al. Bayesian data analysis. 3 edn, (CRC Press, 2014).

44 Lee, M. D. & Wagenmakers, E.-J. in A Practical Course (Cambridge University Press, Cambridge, 2014).

45 Nunez, M. D., Gosai, A., Vandekerckhove, J. & Srinivasan, R. The latency of a visual evoked potential tracks the onset of decision making. Neuroimage 197, 93–108, doi: 10.1016/j.neuroimage.2019.04.052 (2019).

46 Drugowitsch, J., Moreno-Bote, R., Churchland, A. K., Shadlen, M. N. & Pouget, A. The cost of accumulating evidence in perceptual decision making. The Journal of Neuroscience 32, 3612–3628, doi: 10.1523/jneurosci.4010-11.2012 (2012).

47 Lerche, V. & Voss, A. Model complexity in diffusion modeling: benefits of making the model more parsimonious. Front Psychol 7, 1324, doi: 10.3389/fpsyg.2016.01324 (2016).

48 Boehm, U. et al. Estimating across-trial variability parameters of the Diffusion Decision Model: Expert advice and recommendations. Journal of Mathematical Psychology 87, 46–75, doi: 10.1016/j.jmp.2018.09.004 (2018).

49 Heathcote, A., Brown, S. & Mewhort, D. J. Quantile maximum likelihood estimation of response time distributions. Psychon Bull Rev 9, 394–401, doi: 10.3758/bf03196299 (2002).

50 Ditterich, J. Evidence for time-variant decision making. European Journal of Neuroscience 24, 3628–3641, doi: 10.1111/j.1460-9568.2006.05221.x. (2006).

51 Ratcliff, R. et al. Dual diffusion model for single cell recording data from the superior colliculus in a brightness discrimination task. J Neurophysiol 97,(2) 1756-1774. doi.org/10.1152/jn.00393.2006

